# Activity-Induced MeCP2 Phosphorylation Regulates Retinogeniculate Synapse Refinement

**DOI:** 10.1101/2023.07.03.547549

**Authors:** Christopher P. Tzeng, Tess Whitwam, Lisa D. Boxer, Emmy Li, Andrew Silberfeld, Sara Trowbridge, Kevin Mei, Cindy Lin, Rebecca Shamah, Eric C. Griffith, William Renthal, Chinfei Chen, Michael E. Greenberg

## Abstract

Mutations in *MECP2* give rise to Rett syndrome (RTT), an X-linked neurodevelopmental disorder that results in broad cognitive impairments in females. While the exact etiology of RTT symptoms remains unknown, one possible explanation for its clinical presentation is that loss of MeCP2 causes miswiring of neural circuits due to defects in the brain’s capacity to respond to changes in neuronal activity and sensory experience. Here we show that MeCP2 is phosphorylated at four residues in the brain (S86, S274, T308, and S421) in response to neuronal activity, and we generate a quadruple knock-in (QKI) mouse line in which all four activity-dependent sites are mutated to alanines to prevent phosphorylation. QKI mice do not display overt RTT phenotypes or detectable gene expression changes in two brain regions. However, electrophysiological recordings from the retinogeniculate synapse of QKI mice reveal that while synapse elimination is initially normal at P14, it is significantly compromised at P20. Notably, this phenotype is distinct from that previously reported for *Mecp2* null mice, where synapses initially refine but then regress after the third postnatal week. We thus propose a model in which activity-induced phosphorylation of MeCP2 is critical for the proper timing of retinogeniculate synapse maturation specifically during the early postnatal period.

**SIGNIFICANCE STATEMENT:** Rett syndrome (RTT) is an X-linked neurodevelopmental disorder that predominantly affects girls. RTT is caused by loss of function mutations in a single gene MeCP2. Girls with RTT develop normally during their first year of life, but then experience neurological abnormalities including breathing and movement difficulties, loss of speech, and seizures. This study investigates the function of the MeCP2 protein in the brain, and how MeCP2 activity is modulated by sensory experience in early life. Evidence is presented that sensory experience affects MeCP2 function, and that this is required for synaptic pruning in the brain. These findings provide insight into MeCP2 function, and clues as to what goes awry in the brain when the function of MeCP2 is disrupted.

## INTRODUCTION

Rett Syndrome (RTT) is an X-linked neurodevelopmental disorder caused by loss-of-function mutations in methyl-CpG-binding protein 2 (MeCP2), a transcriptional repressor that binds to methylated cytosines across the genome (1, 2). Girls with RTT develop normally during the first year of life, then exhibit developmental stagnation during early childhood, characterized by breathing abnormalities, hand wringing, movement difficulties, seizures, loss of speech, and decelerated head growth (3). While the full etiology underlying the abnormal cognitive and social skills seen in RTT remains unclear, several lines of evidence suggest that synaptic and circuit defects contribute to these symptoms. For example, in mouse models of RTT, abnormalities in various aspects of synaptic development and transmission were observed, including defects in the maturation of excitatory synapses (4), disruption of long-term synaptic plasticity in the hippocampus (5, 6), and altered synaptic scaling (7, 8). Notably, the emergence of RTT symptoms, in both mice and humans, coincides with periods when spontaneous neural activity and sensory experience profoundly influence nervous system development (9, 10). Moreover, mouse models of RTT and MeCP2 duplication syndrome harboring *Mecp2* loss- and gain-of-function mutations, respectively, exhibit a myriad of defects in activity-dependent developmental processes, including dendritic arborization, the maturation of dendritic spines, and synapse elimination (4, 11–13).

Numerous studies in neurons have shown that the MeCP2 protein becomes newly phosphorylated in response to neuronal activity (14–16). Our laboratory and others have identified multiple sites of activity-induced MeCP2 phosphorylation and dephosphorylation, spanning the methyl-DNA-binding domain (MBD), transcriptional repressor domain (TRD), and C-terminal domain (CTD) of MeCP2 (17–22). A variety of effects on MeCP2’s association with methylated DNA and the NCoR co-repressor complex have been ascribed to these post-translational modifications. However, while various *Mecp2* knock-in mouse lines abolishing individual phosphorylation events have been reported to exhibit behavioral and cellular phenotypes (18, 21, 23), a clear understanding of the contribution of these phosphorylation events to the functions of MeCP2 has remained elusive.

Here, we sought to further assess the contribution of activity-induced phosphorylation to MeCP2 function. By mass spectrometry we confirmed the sites of activity-dependent MeCP2 phosphorylation, and then generated a quadruple knock-in (QKI) mouse line in which the four major sites of inducible MeCP2 phosphorylation are mutated to alanines so that the MeCP2 protein is refractory to activity-induced phosphorylation. QKI mice are viable, grossly normal, and exhibit none of the well-characterized RTT phenotypes, making it possible to assess the function of activity-dependent MeCP2 phosphorylation without the confounding effects observed in *Mecp2* null mice. We hypothesized that QKI mice might be specifically defective in steps of activity-dependent brain development, as these would likely be stages when MeCP2 would be subject to activity-induced phosphorylation. To this end, we focused our assessment on the process of early postnatal synapse refinement in the retinogeniculate circuit, as previous studies have shown that the maturation of this circuit is both activity- and MeCP2-dependent (13, 24, 25). We find that, as with *Mecp2* null mice (13), developmental refinement of the synapses between retinal ganglion cells (RGCs) and thalamocortical relay neurons in the dorsal lateral geniculate nucleus (dLGN) is impaired in QKI mice, with more RGC inputs to each relay neuron in QKI mice. Notably, this disruption is observed earlier in QKI mice, between the ages of P14 to 20, compared to *Mecp2* null mice, which show aberrant refinement after P20. Thus, the timing of the synaptic defect in QKI mice suggests that activity-dependent phosphorylation of MeCP2 is critical during the spontaneous-activity dependent phase of retinogeniculate synaptic refinement.

## RESULTS

### Characterization of Activity-Induced Sites of MeCP2 Phosphorylation

We initially sought to assess activity-responsive MeCP2 post-translational modifications, both *in vitro* and *in vivo*, to confirm and extend previous investigations using cultured neurons (18) and epileptic rodent brains (21). To this end, we injected kainic acid into wild-type (WT) mice to induce seizures and then harvested hippocampal tissue after one hour. In parallel, cultured WT mouse cortical neurons were harvested 30 minutes following treatment with either Brain-derived neurotrophic factor (BDNF, 50 ng/mL) or elevated levels of extracellular potassium chloride (KCl, 55 mM) to induce membrane depolarization. In each case, MeCP2 protein was immunoprecipitated and subjected to quantitative mass spectrometry to evaluate possible changes in post-translational modifications.

Consistent with the results of a previous study, which used phosphotryptic mapping of MeCP2 from ^32^P labeled neurons to identify sites of MeCP2 phosphorylation (18), this analysis showed induction of MeCP2 phosphorylation at serine (S) 86, S274, and S421 in response to the various stimuli (**Figure 1A**). Similarly, stimulus-induced phosphorylation of the MeCP2 peptide containing residues threonine (T) 308 and T311 was also observed. Although quantitative mass-spectrometry is unable to distinguish between phosphorylation at these two nearby residues, activity-induced MeCP2 T308 phosphorylation has been previously reported and validated with an MeCP2 phospho-T308-specific antisera (18), strongly suggesting that the detected phospho-peptide species corresponds to phospho-T308. Beyond these four sites, MeCP2 S424 phosphorylation has been previously reported in epileptic mouse brain extracts (21); while we did observe S424 phosphorylation in cultured cortical neurons following KCl-mediated depolarization, this modification was not consistently induced in the hippocampus in response to kainic acid. Likewise, phosphorylation at MeCP2 S80, which has been previously reported to undergo rapid dephosphorylation in response to activity (21), was detected in our dataset, but did not show alterations upon stimulation either *in vitro* or *in vivo*. Finally, this method did not reveal evidence of stimulus-induced changes in other types of MeCP2 post-translational modifications, such as lysine acetylation or arginine methylation.

**Figure 1.**
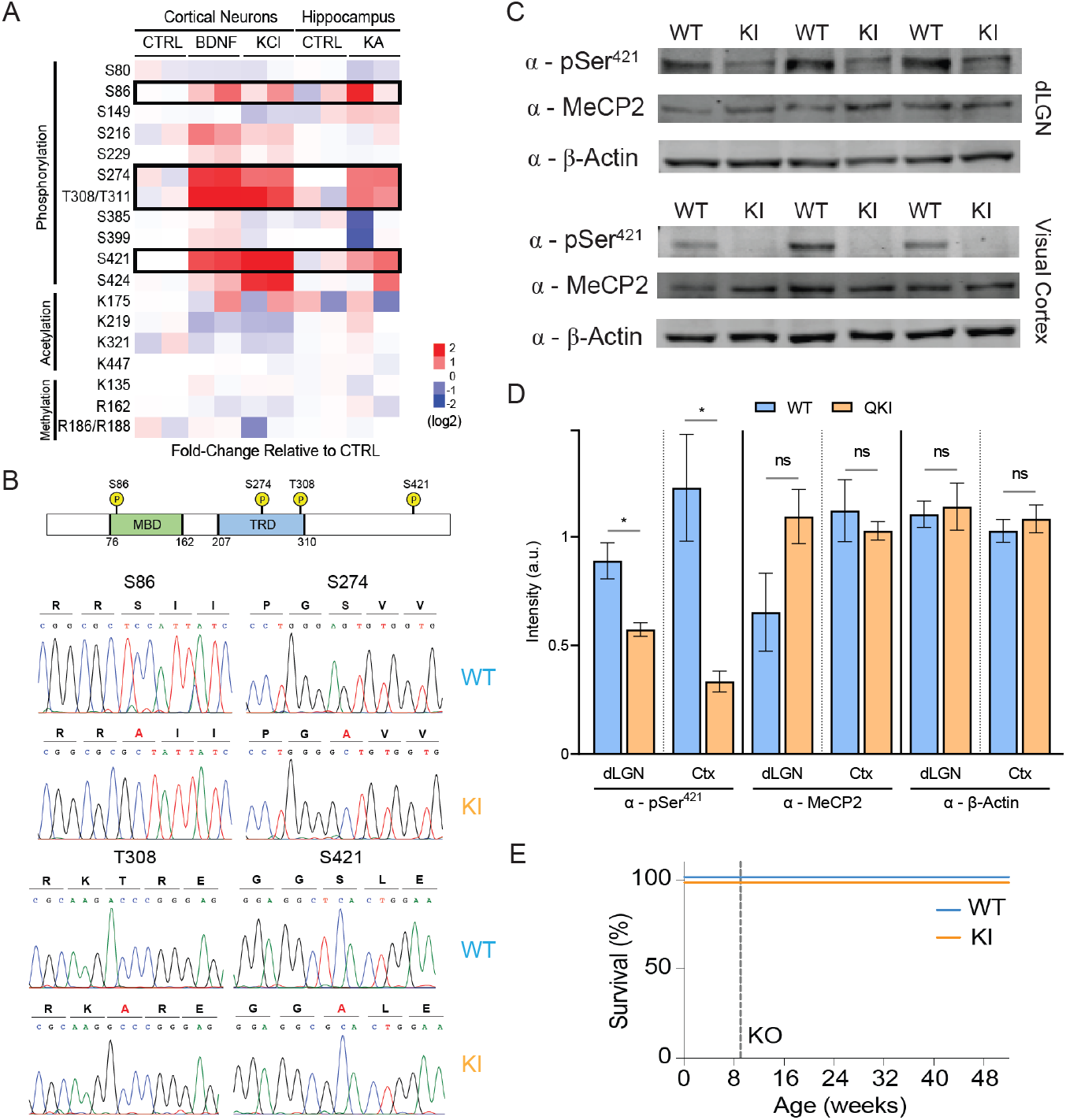
Activity-dependent modifications of MeCP2 and generation of MeCP2 quadruple knock-in (QKI) mice. **A)** Quantitative assessment of stimulus-responsive MeCP2 post-translational modifications. Neuronal activity was induced *in vivo* with kainic acid for 1 hour in mice, after which hippocampi were dissected. Cultured mouse cortical neurons were stimulated at *in vitro* day 7 with BDNF (50 ng/mL) or KCl (55 mM) for 30 minutes to induce depolarization. After induction, MeCP2 protein was isolated using immunoprecipitation, after which samples underwent quantitative mass spectrometry to assess post-translational modifications. Phosphorylation at residues S86, S274, T308/T311, and S421 of MeCP2 were found to be induced by neuronal activity. **B)**Schematic depiction of the MeCP2 protein showing the characterized methyl-DNA-binding domain (MBD) and transcriptional repression domain (TRD), as well as the location of the four sites of activity-responsive phosphorylation targeted for mutation (*Top*). Sanger sequencing confirms the expected base substitutions in QKI mice (*Bottom*). **C)**Western blotting of dLGN and visual cortex from P27-P32 mice with the indicated antisera confirms the loss of MeCP2 S421 phosphorylation in QKI mice without affecting the total levels of MeCP2 protein (n=3 mice). Equal amounts of protein lysates from WT and QKI brain tissue were loaded on a denaturing protein gel, followed by Western blotting with an MeCP2-S421 phospho-specific antisera, an antibody against total MeCP2, and an antibody specific for β-actin as a loading control. **D)**Quantification of Western blots shown in (C). Levels of MeCP2 phosphorylated-S421 are significantly reduced in QKI dLGN and visual cortex compared to WT animals (left). In contrast, levels of total MeCP2 protein are unaffected in QKI mice in both dLGN and visual cortex (right) (n=3 mice). Protein levels in each lane were normalized to the corresponding β-actin control. * p < 0.05, unpaired, two-tailed Student’s t-test. **E)**QKI mice show no differences in longevity as compared to WT littermates. Dotted line indicates typical lifespan on MeCP2 KO mice (51).

While we cannot exclude the presence of other modifications not detected by these methods, our mass spectrometry findings indicate that neuronal stimulation drives the acute phosphorylation of MeCP2 predominantly at four previously reported sites. Intriguingly, three of these sites lie in well-characterized domains important for MeCP2 function—S86 lies in the MBD of MeCP2, and could thus influence MeCP2-chromatin interactions, whereas S274 and T308 both lie in the TRD, where phosphorylation of T308 has been previously shown to disrupt MeCP2 association with the NCoR complex in *in vitro* peptide pulldown experiments (18). While single knock-in mice preventing phosphorylation of S421 or T308 have been generated previously and been shown to have phenotypic differences compared to WT mice (18, 23), the apparent coordinate regulation of these four sites suggests that they may act in concert to modulate neuronal MeCP2 function. We therefore sought to evaluate the effect of mutating these four phosphorylation sites in tandem.

### Generation of MeCP2 QKI Mice

To coordinately abolish activity-induced MeCP2 phosphorylation at S86, S274, T308, and S421, we generated mice in which all four of these sites are converted to alanines (MeCP2 S86A, S274A, T308A, S421A quadruple knock-in, or QKI mice) using standard homologous recombination techniques (23). Sanger sequencing of genomic tail DNA from the resulting animals was used to confirm the presence of the four targeted sequence changes (**Figure 1B**).

Because relatively subtle changes in the level of MeCP2 protein can give rise to significant abnormalities in nervous system development (26), we performed quantitative Western blotting in two distinct brain regions using a total MeCP2 antibody to confirm that MeCP2 protein levels are not significantly altered in QKI mice. Additionally, we used an MeCP2 phospho-S421-specific antisera to confirm that S421 phosphorylation is significantly decreased in both dLGN and visual cortex lysates, with a more pronounced loss of phospho-S421 detected in visual cortex relative to dLGN, perhaps reflecting partial recognition of unphosphorylated MeCP2 in the dLGN by our phospho-S421 antibody (**Figure 1C, D)**. Additionally, the absence of phosphorylation does not significantly affect MeCP2 methyl-DNA binding *in vivo*, as assayed by chromatin immunoprecipitation (ChIP)-sequencing (**Figure S1A**). Consistent with the finding that total MeCP2 protein levels and binding are unchanged, QKI mice are viable, grossly normal, and have normal body weights and lifespans (**Figure 1E, S1B**). Additionally, QKI mice exhibited none of the overt RTT behavioral phenotypes typically seen with decreased MeCP2 protein levels, as judged by an established phenotypic scoring method that measures mobility, gait, hindlimb clasping, tremor, breathing, and general body condition (27) (**Figure S1C**). Thus, the lack of these four phosphorylation events is not sufficient to give rise to pronounced RTT phenotypes, alter MeCP2 protein expression, or affect MeCP2 binding to methyl-DNA.

### Developmental Synaptic Refinement is Aberrant in the dLGN of QKI Mice

While prior studies have implicated MeCP2 in synaptic regulation, the pleiotropic nature of RTT phenotypes has made it difficult to determine if MeCP2 acts as a direct regulator of activity-driven circuit refinement, or whether the synaptic and cognitive deficits observed in RTT mouse models reflect secondary effects of a broader form of neuronal dysfunction. Given the absence of characteristic RTT phenotypes in QKI mice, we went on to assess potential defects in postnatal activity-dependent circuit development in these animals. For these studies, we focused on the retinogeniculate synapses formed between RGCs and thalamocortical (TC) relay neurons of the dLGN, which undergo a well-characterized, sequential process of vision-insensitive (postnatal day (P)0 to P14) and - sensitive (P20 to P30) refinement over the course of early postnatal development (10, 24, 28–31). Importantly, previous work has shown that the process of vision-sensitive synaptic refinement is disrupted in *Mecp2* null mice (13).

To investigate directly the role of activity-induced MeCP2 phosphorylation in this refinement process, we performed whole-cell recordings from TC neurons in acute dLGN slices of QKI and WT littermates at P27-P32 (hereafter referred to as P30 for simplicity), an age after the bulk of synaptic remodeling has occurred, and when there are clear defects in retinogeniculate connectivity in *Mecp2* null mice (13, 24, 25, 32) (**Figure 2**). Following established methods, inward excitatory postsynaptic currents (EPSCs) at -70 mV and outward EPSCs at +40 mV were recorded in a voltage clamp configuration in response to extracellular electrical stimulation of the RGC axons that make up the optic tract. Recordings of currents at maximal, minimal, and intermediate stimulus intensities were made at each holding potential. Particular effort was made to determine the single fiber (SF) response: the amplitude of the EPSC that is first seen following incremental increases in stimulus intensity after failures of synaptic transmission at weaker intensities. We used these values to estimate convergence, or the number of RGCs synapsing onto each TC neuron, using a metric called the fiber fraction (24). The fiber fraction (FF) is the amplitude of the SF EPSC divided by the amplitude of the EPSC in response to maximal stimulus intensity, and thus quantifies the average contribution of each RGC input relative to the total retinal drive to each relay neuron. As such, a lower FF is indicative of more numerous, weak synaptic inputs to each TC neuron, such as those seen early in postnatal development. Previous studies have shown that in WT mice, the FF increases as spontaneous neuronal activity drives refinement of the retinogeniculate synapse from birth until P20, with further increases as the retinogeniculate synapse enters the vision-sensitive period of refinement, indicative of a shift to fewer, stronger RGC inputs (24).

**Figure 2.**
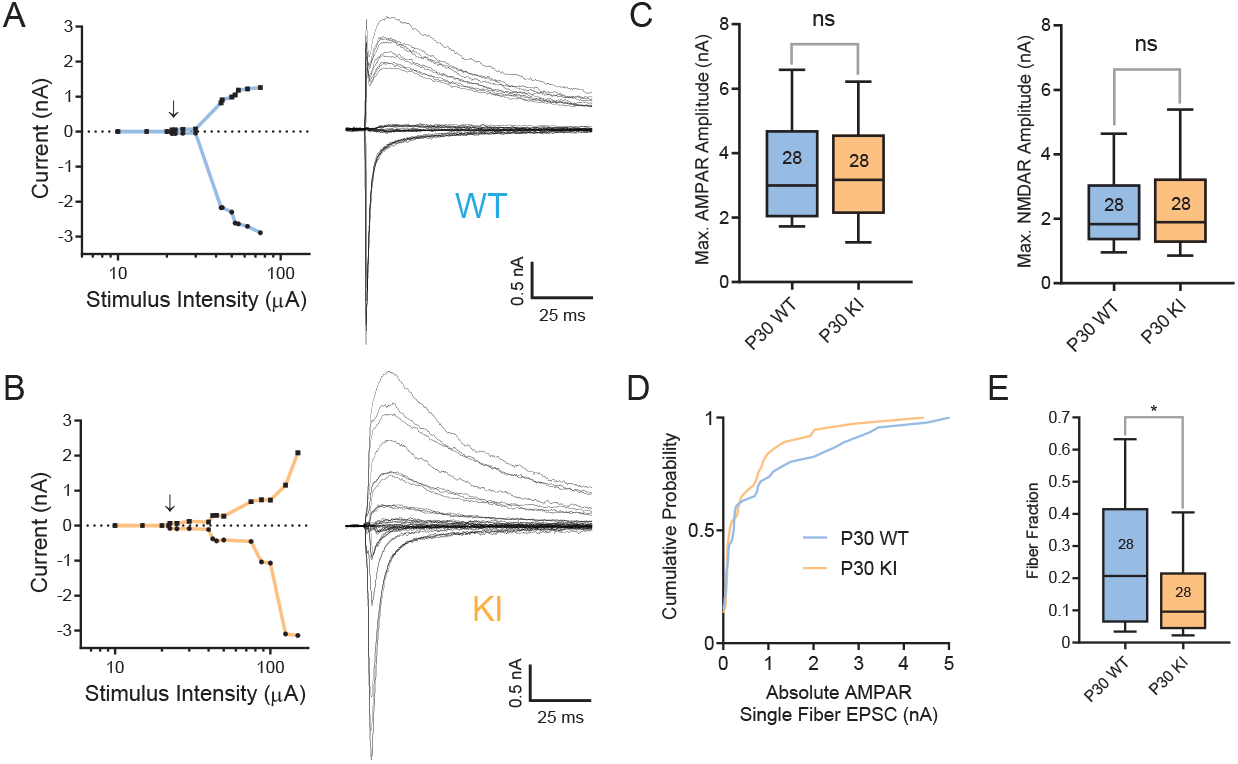
Retinogeniculate synaptic refinement is aberrant in QKI mice. **A-B)** Representative example recordings from P30 WT (A) and QKI (B) relay neurons demonstrating abnormal synaptic refinement in QKI mice. *Left*: Plot of peak AMPAR EPSC (circles) and NMDAR (squares) plotted against stimulus intensity. Arrow indicates the emergence of the first single fiber (SF) following failures of synaptic transmission at lower stimulus intensities. Incremental increases in stimulus intensity were used to sequentially activate additional RGC axons. *Right*: Superimposed representative EPSCs evoked by optic tract stimulation over a range of intensities. At each stimulus intensity, recordings were performed at both -70 mV and +40 mV. Inward currents at -70 mV predominantly represent AMPAR-mediated conductances, whereas outward currents at +40 mV represent AMPAR- and NMDAR-mediated conductances. Stimulus artifacts are blanked for clarity. **C)** Maximal EPSC amplitudes at -70 mV (AMPAR, *Left*) and +40 mV (NMDAR, *Right*) are not significantly different between WT and QKI mice at P30. Numbers in boxplots represent number of cells recorded. **D)** Cumulative probability distributions of AMPAR-mediated SF amplitudes recorded at -70 mV from P30 WT and QKI mice (n=37-46). **E)** The fiber fraction (FF = SF EPSC amplitude/ maximal EPSC amplitude) is significantly reduced in P30 QKI mice.

With incrementally greater stimulus intensities, the peak amplitudes of both the AMPAR and NMDAR EPSCs increase, consistent with the recruitment of more convergent inputs to the recorded cell (**Figure 2A, B**). These recordings suggest a difference between the convergent inputs onto TC neurons in QKI mice versus WT mice. Like the *Mecp2* null mouse, maximal AMPAR- and NMDAR-mediated amplitudes are normal in QKI mice at P30 **(Figure 2C)**. However, unlike *Mecp2* null mice where SF inputs are significantly weaker than those of their WT littermates (13), the cumulative distribution of SF AMPAR amplitudes is not significantly different in QKI mice when compared to WT mice (**Figure 2D**). Nevertheless, examination of the cumulative distributions reveals a clear trend in which a subset of inputs from QKI mice are shifted to the left of WT mice, consistent with fewer inputs strengthening in the LGN of QKI mice. Notably, QKI mice exhibited fewer SF inputs that are greater than 600 pA, a threshold strength that was previously defined as a strong input capable of driving TC neuron firing (29, 33). Since the analysis of maximal currents is not significantly different when QKI and WT mice are compared (**Figure 2C**), the presence of a subset of weaker inputs in QKI mice significantly reduces the median FF (calculated as SF peak amplitude/maximum peak amplitude for each neuron) in QKI mice **(Figure 2E)**. This reduction in FF is consistent with phenotypes previously described in *Mecp2* null mice (13). However, this data suggests that while a subset of TC neurons in QKI mice receives more convergent retinal inputs than TC neurons in WT mice, the overall phenotype of QKI mice is milder than that previously reported of *Mecp2* null mice.

### Retinogeniculate Synapse Refinement is Delayed in QKI Mice

The initial phase of retinogeniculate synaptic refinement, extending from prenatal ages through the second postnatal week, is mediated by spontaneous activity in the form of correlated cholinergic and glutamatergic bursts of action potentials, or waves, from the retina (10, 24, 30, 31, 34). In *Mecp2* null mice, we have previously shown that retinogeniculate refinement proceeds relatively normally until P20, after which connectivity regresses (13). Between P20-P30, the number of convergent inputs increases, while the average strength of the inputs decreases in *Mecp2* null mice compared to WT mice. We thus asked whether a similar time course of disrupted refinement is seen in QKI mice. Given previous findings that MeCP2 S421 phosphorylation is present in the visual cortex around the time of eye-opening (23), we first investigated whether early-stage refinement at P14-P15 (hereafter referred to as P14 for simplicity) is disrupted in QKI mice. In this regard, whole-cell recordings from TC neurons in acute P14 dLGN slices prepared from P14 QKI and WT littermates revealed no differences in single fiber AMPAR- and NMDAR-mediated EPSC amplitudes, maximal EPSC amplitudes, or FF between the two genotypes (**Figure 3A *Left*, 3C, 3D**). Thus, mimicking trends previously observed in *Mecp2* null mice (13), the early spontaneous activity-driven period of refinement is unaffected by the loss of activity-induced MeCP2 phosphorylation, with both WT and QKI mice exhibiting a predominance of weak RGC synapses onto each TC neuron around eye-opening.

**Figure 3.**
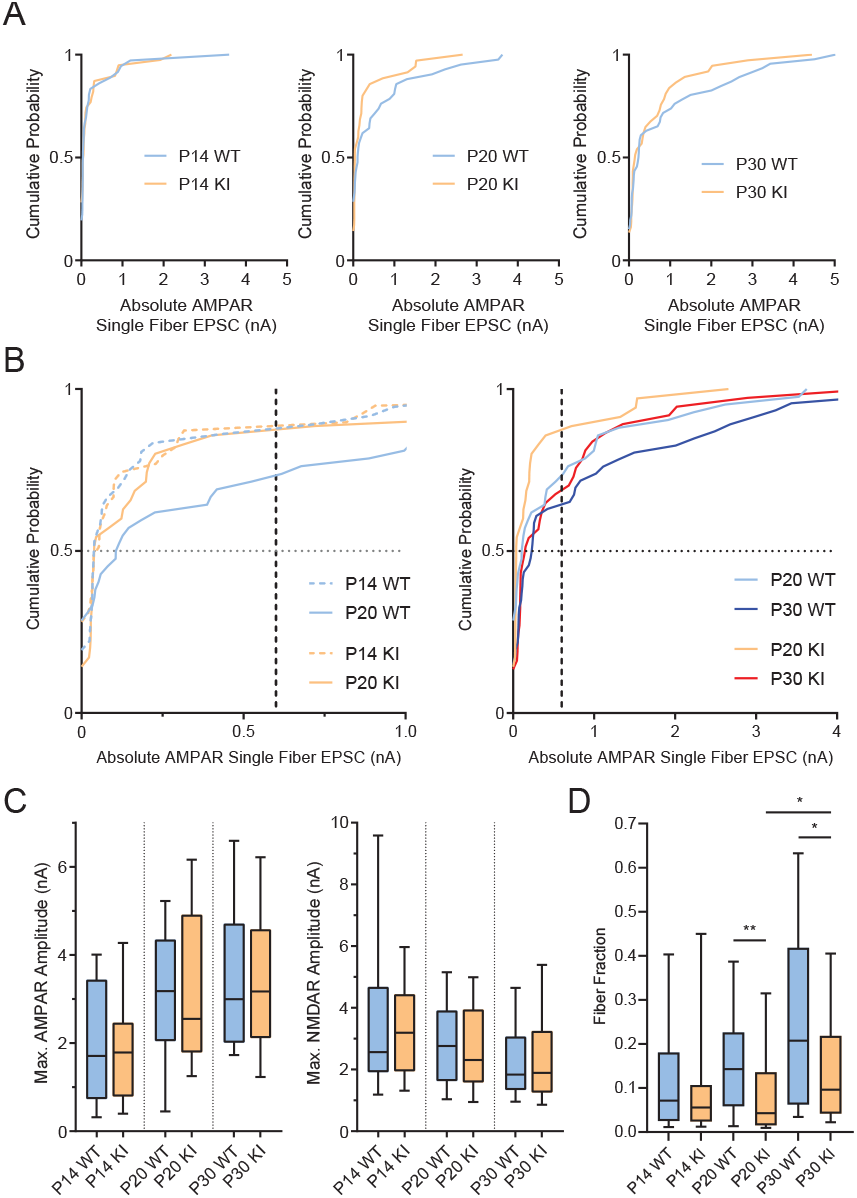
Delayed retinogeniculate refinement in QKI mice. **A)** Cumulative probability distributions of AMPAR-mediated single fiber (SF) amplitudes at -70 mV for P14 (*Left*), P20 (*Middle*) and P30 (*Right*) QKI and WT mice. SF AMPAR EPSC amplitude distributions of WT and QKI mice overlap at P14. By P20, a subset of inputs in WT mice have strengthened, shifting the distribution to the right compared to that of QKI (WT vs. QKI: 3rd & 4th quartiles p<0.02; full distribution p =0.5). By P30, the QKI distribution shifts toward that of the WT, indicating some strengthening of the SF inputs in the mutant (WT vs. QKI: 3rd & 4th quartiles p=0.16; full distribution p=0.64). **B)** Comparison of cumulative probability distributions of AMPAR-mediated SF amplitudes (from panel A) between P14 vs. P20 (*Left*), and P20 vs. P30 (*Right*), from WT and QKI mice. The median (dotted horizontal line at y=0.5) WT SF strengthens by 187% (0.039 at P14 to 0.11 at P20) whereas the median QKI SF slightly weakens by -27% (0.054 at P14 to 0.039 at P20). AMPAR-mediated SFs begin to strengthen from P20 to P30 in the QKI (*Right*), visualized by the median values at y=0.5 in the WT (0.11 at P20 to 0.23 at P30, a 109% increase) versus in the QKI (0.039 at P20 to 0.15 at P30, a 285% increase). Dashed vertical lines at x=0.6 nA highlight the greater proportion of weak RGC inputs less than 0.6 nA in the QKI relative to WT at P20. By P30, the proportion of weak RGC inputs less than 0.6 nA is relatively closer in value between QKI and WT mice. **C)** Maximal EPSC amplitudes at -70 mV (AMPAR, *Left*) and +40 mV (NMDAR, *Right*) are not significantly different between WT and QKI mice at all ages tested. **D)** The FF is not significantly different between WT and QKI mice at P14. By P20, the FF is significantly lower, indicating the presence of more RGC inputs to each relay neuron in QKI mice. While the FF of QKI mice increases significantly between P20 and P30, it does not recover to WT levels. All statistics were performed using a two-tailed Mann-Whitney U, Kolmogorov-Smirnov, or Kruskal-Wallis when comparing more than two groups. ns, p > 0.05; *, p < 0.05; **, p < 0.01. Box, 25%–75% interquartile range; whiskers, 10%–90% interquartile range.

Notably, however, we found that the time course of disrupted refinement in the QKI mice is different from that previously described for *Mecp2* null mice. We performed recordings of P19-P21 WT and QKI mice (hereafter referred to as P20 for simplicity) and found that the FF does not significantly increase between P14 and P20 in QKI mice when comparing synaptic connectivity across three age groups (P14, P20, and P30). First, comparison of the cumulative distribution of SF peak amplitudes suggests that while WT mice exhibit a significant increase in SF peak amplitudes between P14 and P20, very few RGC inputs strengthen in QKI mice over the same developmental window (**Figure 3A, 3B *Left***). Notably, the median SF amplitude at -70 mV strengthens by 187% in WT mice from P14 to P20 (0.039 to 0.11), whereas the median SF amplitude weakens by 27% in QKI mice over this period (0.054 to 0.039). Quantification of the third and fourth quartiles of the cumulative distributions, which represent the subset of inputs that have strengthened, reveals a significant difference between WT and QKI mice at P20 (p<0.02, Mann-Whitney). This is in contrast to *Mecp2* null mice, in which the disruption of refinement does not occur until after P20 (13). Strikingly, between P20 and P30, SF inputs of QKI mice significantly strengthen (P20 vs P30, p=0.005, Kruskal-Wallis, n=35-37 cells), so that they are no longer significantly different in strength than the SF inputs of WT mice (WT vs QKI mice, 3^rd^ and 4^th^ quartiles, p=0.16) (**Figure 3B, *Left vs. Right panels***).

We then compared maximal currents and the FF across development of WT and QKI mice. Maximal peak AMPAR and NMDAR EPSC currents were not significantly different at any point during development **(Figure 3C)**; however, the FF shows distinct developmental trends between the two genotypes **(Figure 3D)**. The median FF in WT mice doubles between P14 and P20 (0.07 to 0.14), consistent with the pruning of convergent retinal inputs, while the median FF in QKI mice does not significantly change over this same period (0.06 to 0.04). Instead, the FF in QKI mice only increases between P20 vs P30 (p=0.005, Kolmogorov-Smirnov test, n=35-37), although not to the same level as in WT mice (**Figure 3D**). Taken together, retinogeniculate refinement appears to be delayed in QKI mice at P20, but partially recovers by P30.

We next examined other synaptic features that are known to change during postnatal retinogeniculate development (24, 28). We found no significant differences in the AMPAR/NMDAR ratio (calculated as the ratio of the maximal AMPAR-mediated EPSC at -70 mV divided by the maximal NMDAR-mediated EPSC at +40 mV) when WT and QKI mice are compared. However, there are notable differences in the EPSC waveform when WT and QKI mice are compared—EPSCs recorded from P20 QKI mice at both holding potentials exhibit significantly slower decay kinetics (t) than those of WT mice at the same age. By P30, EPSC decay kinetics have accelerated in QKI mice so that the AMPAR and NMDAR decay kinetics are similar to those seen in WT mice. Overall, this suggests that the development of the TC synapse in QKI mice is delayed relative to WT mice (28) (**Figure 4A**).

**Figure 4.**
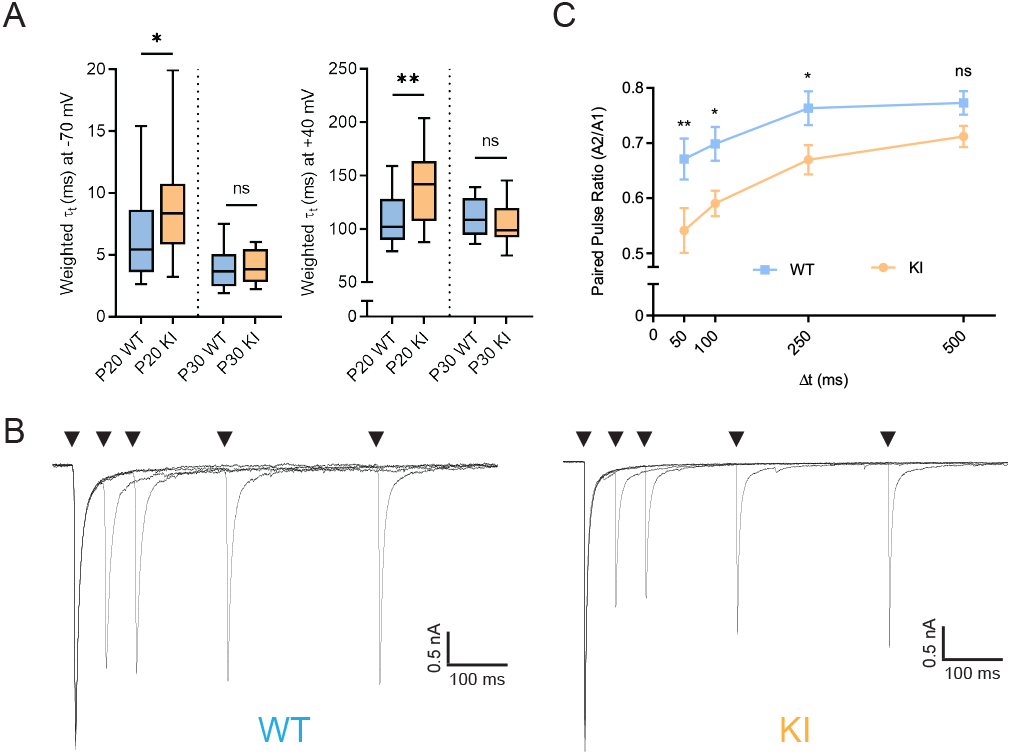
Retinogeniculate synaptic properties are altered in QKI mice. A) Decay kinetics of EPSCs at -70 mV (*Left*) and +40 mV (*Right*) are significantly slower in QKI at P20, but not at P30, as compared to WT mice. B) Superimposed pairs of EPSCs recorded at a holding potential of -70 mV with varying ISIs (50,100, 250, and 500 ms) from P29-32 WT (*Left*) and QKI (*Right*) mice. Stimulus artifacts were blanked for clarity. C) Mean paired pulse ratio (PPR) ± standard error of the mean, calculated by dividing the amplitude of the second EPSC (A2) by the amplitude of the first EPSC (A1), shows greater short-term depression in QKI mice as compared to WT mice at all inter-stimulus intervals (ISIs) tested. WT: n=17 from 3 mice; KI: n=14 cells from 4 mice. Statistics in (A) were performed using a two-tailed Mann-Whitney U test, and those in (B) using a one-way ANOVA and Fisher’s LSD of multiple comparisons. ns, p > 0.05; *, p < 0.05; **, p < 0.01.

### Retinogeniculate Synapse Refinement Defects in QKI Mice are Presynaptic

One possible explanation for the presence of weaker SF inputs in QKI mice is that the probability of vesicle release from RGCs is decreased in QKI animals. Towards this end, we stimulated the optic tract twice in rapid succession with varying inter-stimulus intervals (ISIs) to determine paired-pulse ratios (PPRs) in P29-32 WT and QKI mice. PPRs were found to be significantly decreased in QKI mice compared to their WT littermates across a range of ISIs (**Figure 4B, 4C**), indicating that vesicles in QKI mice are depleted faster and/or recover more slowly than in WT mice. Importantly, *Mecp2* null mice have been previously reported to not exhibit alterations in PPR at P30, and prior studies have shown that deficits in synaptic refinement do not necessarily lead to changes in PPRs (13, 35). Our findings thus suggest that activity-induced MeCP2 phosphorylation contributes to retinogeniculate synapse refinement at least in part by modulating pre-synaptic vesicle release in RGCs.

### RGC axon targeting to dLGN is not disrupted in QKI mice

The deficits observed in synaptic refinement in QKI mice at P20 led us to hypothesize that these effects might be due to defects in the structural development of the retinogeniculate synapse. To investigate this possibility, we examined eye-specific RGC axon segregation in QKI mice. While the segregation of retinal axons into eye-specific domains is largely complete by P8-P10 in WT mice (13, 36), a failure to maintain this organization has been previously observed in *Mecp2* null animals when assessed later in development (P46-P51) (13). To determine if eye-specific segregation is altered in QKI mice, we injected cholera toxin subunit B conjugated to different fluorescent dyes into each eye of littermate WT and QKI mice at P58-P64 to visualize retinal projections to the dLGN (**Figure S2)**. Segregation was quantified by calculating the logarithm of the ratio of ipsilateral to contralateral intensity for each fluorophore at every pixel, known as the R-value. The mean variance of R from each section was then used to assess segregation, where a high value indicates a high degree of eye-specific segregation and a low value suggests a low degree of segregation (37). Surprisingly, in contrast to the significant defect in eye-specific segregation observed in *Mecp2* null mice, this analysis revealed no significant differences in eye-specific segregation between QKI and WT mice (**Figure S2C**). These results are consistent with the idea that, in contrast to *Mecp2* null mice, retinogeniculate circuits in QKI mice do not regress with age.

### No Detectable Changes in Gene Expression in Visual Circuits of QKI Mice

Given the well-characterized role of MeCP2 as a transcriptional repressor, we next sought to characterize transcriptional changes in QKI mice that might give rise to the observed electrophysiological abnormalities. As three of the activity-dependent phosphorylation sites of MeCP2 occur within its MBD and TRD, one possibility is that activity-dependent MeCP2 phosphorylation relieves this repressor function by inhibiting the binding of MeCP2 to methylated DNA and/or inhibiting the interaction of MeCP2 with NCoR, thus allowing for or facilitating the induction of activity-dependent genes. In the absence of activity-dependent phosphorylation of MeCP2 in QKI mice, the failure to relieve MeCP2 repression might thus result in a defect in the induction of activity-dependent genes. To begin to investigate this possibility, we used bulk RNA-seq to profile gene expression in dLGN and visual cortex of WT and QKI mice at P16 (n=10 per genotype) and P15 (n=6 per genotype), around the time of eye-opening but before the onset of vision-sensitive synaptic refinement, as we hypothesized that gene expression changes would precede the observed synaptic refinement defects observed at P20. However, differential gene expression analysis identified no significant gene expression changes between WT and QKI mice at these early developmental stages, or later during the vision-sensitive period when we assessed gene expression at P30 (n=6 per genotype) (**Figure 5A, S5)**. One possible explanation for these findings is that alterations in gene expression in QKI mice are subtle, especially given that overall gene expression fold-changes in *Mecp2* null mice, while encompassing many genes, are quite modest and thus may be below the level of detection in QKI mice.

**Figure 5.**
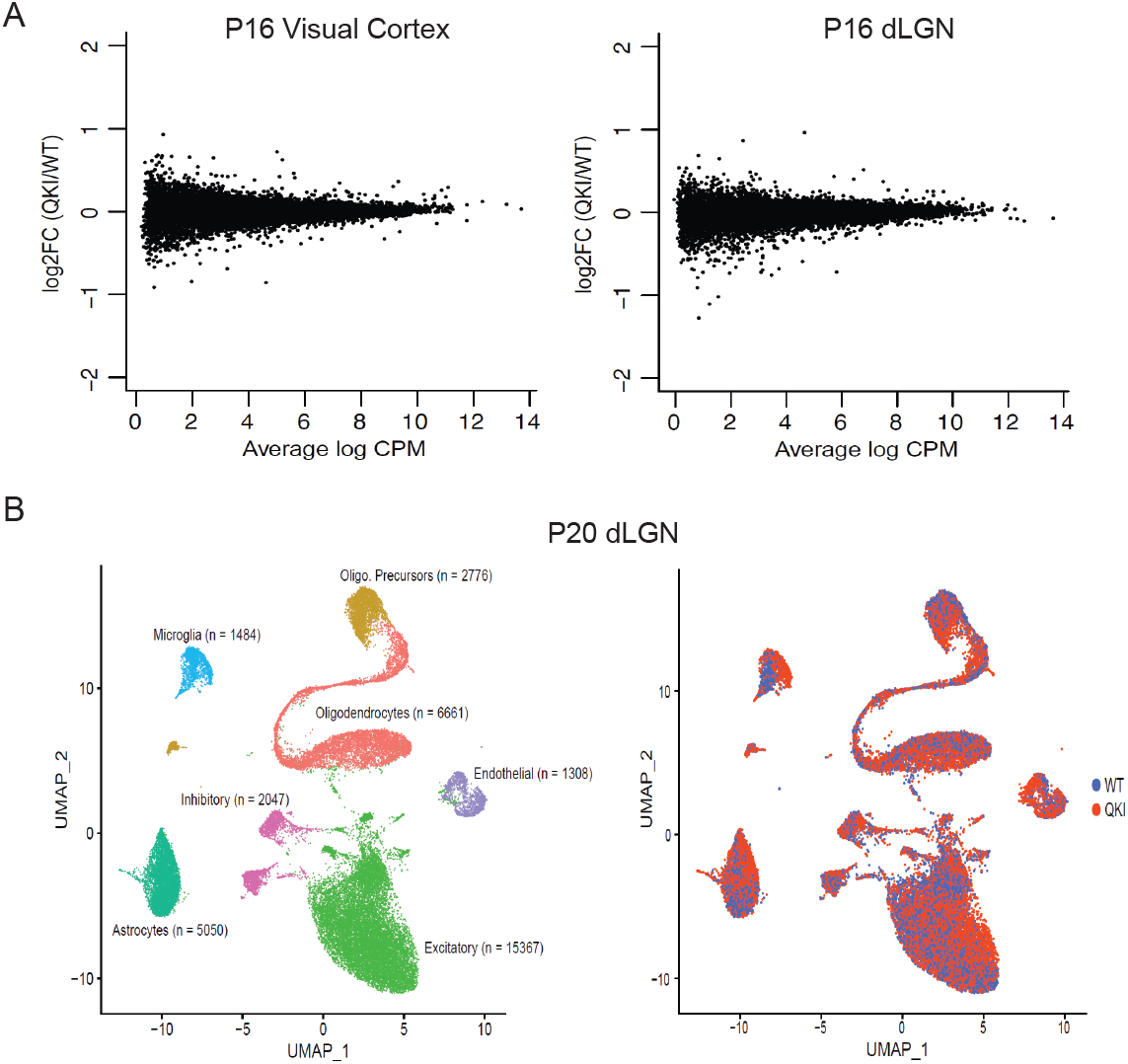
No detectable gene dysregulation in quadruple knock-in (QKI) mice. **A)** Mean-difference plot of differential expression analysis from bulk RNA-sequencing of 10 WT and 10 QKI visual cortex (*Left*) and dLGN (*Right*) at P16. No significant differentially expressed genes were detected. Significance P>0.05. Fisher’s exact test with Benjamini-Hochberg correction. **B)** UMAP visualization of WT and QKI snRNA-seq data from P20 dLGN revealed no differentially expressed genes between WT and QKI mice in excitatory neurons, inhibitory neurons, astrocytes, oligodendrocytes and oligodendrocyte precursors, or endothelial cells. One differentially expressed gene, *Atg16l2*, was detected in the microglia cluster. Three mice were pooled per genotype (n=2 replicates, 12 mice total). Cell types were assigned by known marker genes as discussed in Methods. Nuclei are colored by assigned cell type (*Left*) or by genotype (*Right*). n=number of nuclei per cluster. Oligo., oligodendrocytes.

Given these results and the neuronal activity-dependence of these MeCP2 phosphorylation events, we also assessed vision-dependent gene expression by dark rearing mice from P20-P30 (known as late dark rearing (LDR)) and then acutely re-exposing the mice to light for either zero or four hours (35), reasoning that the effects of disrupting MeCP2 phosphorylation on gene expression might be easier to detect upon deprivation and subsequent synchronization of visual experience. To validate our stimulation protocol, we first performed quantitative reverse transcription PCR (RT-qPCR) of RNA extracted from both visual cortex and dLGN using primers designed to anneal to the RNA transcripts of known activity-regulated genes, including the immediate early proto-oncogene *Fos* (38, 39), the bHLH-PAS transcription factor *Npas4* (40–42), brain-derived neurotrophic factor *Bdnf* (43–45), and the cytokine receptor *Tnfrsf12a* / *Fn14* (35). Following LDR and four hours of light exposure, we indeed detected a significant induction of these genes either in dLGN only (*Fn14*), or both brain regions (*Fos, Npas4, Bdnf*) using both RT-qPCR and bulk RNA-seq, as previously reported (**Figure S3)**. The successful induction of known activity-regulated genes enabled us to use this LDR and light exposure paradigm to assess gene expression in response to neuronal activity in both the dLGN and visual cortex of WT and QKI mice. No significant gene expression differences were observed between WT and QKI samples from either dLGN or visual cortex at both zero and four hours of light stimulation after LDR (**Figure S4)**. However, given that the synaptic phenotype of QKI mice is recovering between P20-P30, differences between LDR QKI and WT mice may be obscured.

Reasoning that bulk RNA-seq might not be sensitive enough to detect cell type-specific differences in gene expression between WT and QKI mice, we also performed single-nucleus RNA-seq of dLGN from WT and QKI mice. Due to the small size of the dLGN, tissue from three P20 animals was pooled for each biological replicate (n=2 replicates, 6 mice per genotype). After nuclei collection, single-cell libraries were generated using 10x Genomics, followed by data analysis with the 10x Genomics pipeline and Seurat (46). Cell types were subsequently assigned on the basis of known marker genes, including *Gad2* for inhibitory neurons, *Aldoc* for astrocytes, and *Slc17a7* for excitatory neurons (47). Clusters expressing the same marker genes were then combined, after which differential gene expression analysis between WT and QKI mice was run within each cluster and across all cells (a total of 34,693 nuclei were included in final analysis) (**Figure 5B, S5C, S5D**). Across all cells, this analysis identified no differentially expressed genes between WT and QKI mice. Furthermore, we saw no differentially expressed genes between WT and QKI mice within excitatory neurons, inhibitory neurons, astrocytes, oligodendrocytes and oligodendrocyte precursors, or endothelial cells. Within the microglia cluster, we detected only one differentially expressed gene, *Atg16l2*. While the lack of differentially expressed genes is surprising, it is possible that any differentially expressed genes between WT and QKI mice are below the level of current detection, that the detected synaptic changes are due to gene expression alterations that occur presynaptically within RGCs, or that the synaptic refinement defects are being driven by a function of MeCP2 separate from its role as a transcriptional repressor.

## DISCUSSION

Previous studies have suggested a role for MeCP2 in both spontaneous activity-dependent and experience-driven synaptic regulation. For example, in addition to deficits in experience-dependent retinogeniculate synapse remodeling in *Mecp2* null mice (13), MeCP2 loss has also been reported to accelerate the critical period of ocular dominance plasticity in visual cortex and to disrupt vision- and activity-dependent homeostatic synaptic scaling responses (7, 8, 48). However, it has been difficult to distinguish the direct contribution of MeCP2 to activity-driven processes from the secondary effects stemming from the generalized RTT-associated neural dysfunction concurrent with MeCP2 loss-of-function models. To begin to address this issue, we generated a novel mouse line in which MeCP2 is no longer susceptible to activity-induced phosphorylation. While these animals appear grossly normal and fail to exhibit hallmark symptoms of RTT, they have deficits in retinogeniculate circuit refinement.

Intriguingly, QKI mice do not phenocopy the significant deficits in retinogeniculate refinement reported previously in *Mecp2* null mice (13). Rather than presenting with a milder form or a subset of the characteristics of *Mecp2* null mice, the QKI mice exhibit a refinement defect that is distinct from that of *Mecp2* null mice. Importantly, the timing of developmental retinogeniculate refinement of WT littermates of QKI mice is remarkably similar to that reported previously for WT littermates of *Mecp2* null mice (13), justifying the validity of this comparison between these two mutants. Despite the fact that the two mutants exhibit similar aberrant retinogeniculate connectivity at P30, the phenotype of each of the mutants is a consequence of different developmental trajectories. In *Mecp2* null mice, refinement is not significantly different from WT littermates during the vision-independent phase of development (from birth to P20), after which the retinogeniculate circuit regresses as the number of convergent RGC inputs increases while the average strength of the inputs decreases. In contrast, synaptic connectivity in QKI mice appears normal at P14, after which further synapse elimination and strengthening is disrupted (**Figure 3B left, 3D**). That is, QKI mice exhibit disrupted synaptic refinement after P14, whereas this disruption occurs later, after P20, in *Mecp2* null mice. By P20, when compared to WT littermates, the strength of individual RGC axons is weaker, decay kinetics of EPSCs are slower, and RGC inputs fail to eliminate correctly in QKI mice. Notably, in QKI mice, it is only after P20 that there is further refinement, with significant strengthening of SF inputs, a decrease in convergent inputs, and acceleration of EPSC decay kinetics. However, synaptic refinement fails to fully normalize by P30, as the number of convergent inputs onto TC neurons is still significantly greater in QKI mice when compared to their WT littermates. Together, these findings provide strong support for a direct role for activity-regulated MeCP2 phosphorylation in activity-dependent synaptic remodeling prior to the vision sensitive period of synaptic refinement.

Our findings are most consistent with a scenario where phosphorylation of the four activity-dependent sites on MeCP2 is necessary for the proper timing of retinogeniculate development. The delayed acceleration of EPSC kinetics is also consistent with this model. It is possible that the functional difference between the QKI and WT mice diminishes further with age; however, recordings at later time points are difficult to obtain due to technical limitations. The fact that eye-specific segregation is indistinguishable between WT and QKI adult mice at a later age further suggests that, while initially delayed in QKI mice, the degree of refinement of the retinogeniculate synapse eventually begins to recover towards the level observed in WT mice. The distinct nature of the refinement defect observed in QKI mice compared to *Mecp2* null animals is notable, and could reflect a negative regulatory role of these phosphorylation events with respect to MeCP2 function. Consistent with this idea, previous work from Ebert et al. (2013) suggests that MeCP2 T308 phosphorylation suppresses MeCP2’s ability to repress transcription. Loss of phosphorylation could thus reflect enhanced MeCP2 activity in some contexts, which may also account for why the vision-insensitive phase of retinogeniculate refinement is disrupted in QKI mice but not in *Mecp2* null mice. In this regard, MeCP2 function could be largely inhibited by phosphorylation during the early phase of synaptic refinement, thereby minimizing the effects of MeCP2 deletion at this stage.

Despite the known role of MeCP2 as a transcriptional repressor, we were surprised to find that loss of activity-dependent phosphorylation of MeCP2 does not have a detectable impact on gene expression in the dLGN or visual cortex at the time points assessed, raising the possibility that other mechanisms could drive the differences seen in synaptic refinement between WT and QKI mice. Alternatively, alterations in gene expression that underlie the synaptic defects in QKI mice may simply be too subtle to detect with current methods. This idea is consistent with the relatively small fold-changes seen in genes when comparing WT and *Mecp2* null mice (49).

Nevertheless, it is also possible that presynaptic mechanisms could account for the failed refinement in QKI mice, such as disrupted feedforward excitation from RGCs, in line with our PPR studies showing that vesicle release probability is altered in RGCs in QKI mice. Thus, gene expression changes in RGCs, rather than in the dLGN, could be responsible for the synaptic defects that we have observed in QKI mice.

The absence of MeCP2 phosphorylation in QKI mice might still have been expected to affect gene expression in the dLGN given previous reports that MeCP2 phosphorylation disrupts the association of MeCP2 with the NCoR corepressor complex *in vitro*. However, while attenuated light-dependent gene responses have been previously reported in the visual cortex of MeCP2 T308A knock-in mice (18), these effects were not seen after substantial backcrossing of the mice (unpublished observation). Thus, whether the loss of activity-induced MeCP2 phosphorylation affects circuit refinement by subtly affecting gene expression or by some other mechanism remains to be determined.

Beyond the retinogeniculate system, further study of the QKI mice will be needed to address the broader contribution of MeCP2 phosphorylation to postnatal experience-dependent circuit remodeling. In this regard, MeCP2 has been implicated in the control of ocular dominance plasticity, as well as the timing of critical period plasticity (48, 50). Likewise, it will also be of interest to examine potential deficits in neuroplasticity in adult QKI mice. Ultimately, it will be important to dissect the contribution of individual MeCP2 phosphorylation sites to these phenotypes through the assessment of synaptic refinement in single knock-in MeCP2 phospho-mutant mouse strains, as well as to elucidate the mechanisms through which activity-dependent phosphorylation of MeCP2 affects synaptic refinement.

## MATERIALS AND METHODS

### Generation of Mecp2 Quadruple knock-in mice

All animal experiments were approved by the National Institutes of Health and the Harvard Medical School Institutional Animal Care and Use Committee and were conducted in compliance with the relevant ethical regulations. MeCP2 quadruple knock-in mice were generated by homologous recombination in embryonic stem (ES) cells as described previously (18, 23) with minor modifications. The *Mecp2* targeting construct generated previously (23) contains a floxed neomycin-resistance cassette (NEO) in *Mecp2* intron 3 and 4.1 kb of the *Mecp2* locus upstream (including exon 3, which contains S86) and 3.9 kb downstream (including *Mecp2* exon 4, which contains S274, T308, and S421). MeCP2 S86A, S274A, T308A, and S421A mutations were added to this targeting construct, and the sequence of the entire targeting construct was verified by Sanger sequencing prior to use in gene targeting. Gene targeting was performed at the IDDRC Mouse Gene Manipulation Core at Boston Children’s Hospital. The construct was linearized by digesting with NotI and electroporated into JM8 Agouti C57BL/6N mouse ES cells. Genomic DNA was isolated from G418-resistant ES cell clones and was screened by PCR for each arm of the targeting construct. One primer recognized a sequence located outside the targeted region, while the other primer recognized a sequence located in the other arm of the targeting construct. A 1.8 kb increase in the size of the product indicated successful insertion of the NEO cassette into the *Mecp2* locus. Two confirmed MeCP2 S86A; S274A; T308A; S421A ES cell clones were transfected using a Cre recombinase-expressing plasmid. Appropriate excision of the loxP-flanked NEO cassette was subsequently confirmed by PCR genotyping, and the presence of the four mutations was verified by PCR and Sanger sequencing. The MeCP2 QKI ES cell clones were injected into C57BL/6 blastocysts and subsequently implanted into pseudopregnant females. The resulting chimeric offspring were mated with C57BL/6J mice, and the offspring were screened by PCR genotyping of tail genomic DNA for the presence of the residual loxP site to confirm germline transmission of the mutant allele. The sequence of all four mutations was verified by Sanger sequencing of tail genomic DNA. Mice were housed under a standard 12-hour light cycle unless otherwise indicated (see late dark rear methods).

### Quantitative mass spectrometry for MeCP2 modifications

#### Induction of neuronal activity

MeCP2 immunoprecipitation for quantitative mass spectrometry was performed from both cultured embryonic mouse cortical neurons and adult mouse hippocampus. For cultured cortical neurons, embryonic day 16.5 (E16.5) cortices from wild-type C57BL/6 mice were dissociated with Papain (Sigma). Neurons were plated at 30 million cells in 15 cm dishes pre-coated overnight with 40 *μ*g/mL poly-D-lysine and 4 *μ*g/mL laminin. Neurons were grown in Neurobasal media (Gibco) containing 1X B-27 Supplement, 1X GlutaMAX supplement, and 1X penicillin-streptomycin. After 6 days in vitro (DIV6), neurons were silenced for 16 hours with 1 *μ*M TTX and 100 *μ*M AP5 to prevent spurious activity. On DIV7, neurons were either left untreated (control) or stimulated for 30 minutes by depolarizing with 55 mM potassium chloride (KCl) or by adding 50 ng/mL of Brain-derived neurotrophic factor (BDNF). For each condition, 180 million neurons were used (untreated, KCl, BDNF) and two biological replicates were performed. After 30 minutes, cells were harvested by washing once with cold PBS, then scraped in 4 mL cold NE1 buffer per 30 million cells (NE1: 20 mM HEPES pH 7.9, 10 mM KCl, 1 mM MgCl_2_, 0.1% Triton X-100, 1 mM DTT, 1X Protease Inhibitor Cocktail (Roche), 1% Phosphatase Inhibitor Cocktail 2 and 3 (Sigma), and 10 mM sodium butyrate). For adult mouse hippocampus, 8-week-old male wild-type C57BL/6 mice were injected intraperitoneally with 15 mg/kg kainic acid. Mice were sacrificed 1 hour after injection, and the hippocampus was dissected. Hippocampi from 6 mice were combined for each control and kainic acid condition, and two biological replicates were performed. Hippocampi from a single brain were homogenized with a pellet pestle homogenizer in 50 *μ*L NE1 buffer, then the volume was increased to 4 mL NE1 for cell lysis.

#### MeCP2 immunoprecipitation

For both cultured cortical neurons and hippocampal tissue, cell lysis was ensured by pipetting up and down 10 times and rotating for 10 minutes at 4°C. Nuclei were pelleted by spinning at 500 x g for 10 minutes at 4°C. Nuclei were resuspended in one pellet volume of NE1 buffer, and 0.9 *μ*l Benzonase (Sigma) per 90 million cells or hippocampi from 3 brains was added, and rotated for 30 minutes at 4°C. Benzonase activity was quenched by adding 5.67 volumes of NE1 250 (NE1 buffer with 250 mM NaCl and 0.5% Triton X-100) and rotating for 20 minutes at 4°C. Insoluble material was removed by spinning at 16,000 x g for 20 minutes at 4°C. The nuclear supernatant was pre-cleared by adding 30 *μ*L Protein A Dynabeads (Invitrogen) per 90 million neurons or hippocampi from 3 brains and rotating for 30 minutes at 4°C. MeCP2 antibody (14) was conjugated to Protein A Dynabeads in NE1 250 by rotating at 4°C for 1.5 hours. Antibody-coupled beads were then added to pre-cleared nuclear supernatant, and immunoprecipitation was performed by rotating tubes for 1.5 hours at 4°C (for 90 million cells or hippocampi from 3 brains, 90 *μ*L of MeCP2 antibody with 270 *μ*L beads were used). After immunoprecipitation, beads were washed 4 times for 5 minutes each in NE1 250 with rotation at 4°C. Proteins were eluted from beads by adding 1X sample buffer (1X LDS (Life Technologies) in PBS with 5% Beta-Mercaptoethanol) and boiling for 15 minutes at 95°C. 10% of eluted sample was used to verify successful MeCP2 IP by Western blot, and 90% of supernatant was electrophoresed on a 10% Bis-Tris gel (Invitrogen) and stained with QC Colloidal Coomassie (Bio-Rad). The MeCP2 bands were excised using clean razor blades and submitted for mass spectrometry.

#### TMT Quantitative Mass Spectrometry and Analysis

Mass spectrometry was performed at the Thermo Fisher Center for Multiplexed Proteomics at Harvard Medical School. Bands were de-stained, reduced with DTT, and alkylated before digesting overnight with Trypsin. Peptides were extracted from gels, dried, and desalted. Cleaned peptides were labeled with TMT10 reagents. Labeling reactions were combined, cleaned, and dried down. Peptides were resuspended in 5% Acetonitrile, 5% formic acid, and the entire sample was analyzed on an Orbitrap Fusion Mass spectrometer. Peptides were separated using a gradient of 6 to 28% acetonitrile in 0.125% formic acid over 180 minutes. Peptides were detected (MS1) and quantified (MS3) in the Orbitrap. Peptides were sequenced (MS2) in the ion trap. MS2 spectra were searched using the SEQUEST algorithm against a Uniprot composite database derived from the mouse proteome containing its reversed complements and known contaminants. MeCP2 was confirmed as the top protein in the sample, and then raw files were searched with a composite database including only the MeCP2 sequence, contaminants, and reverse complements to identify modifications of interest. Peptide spectral matches were filtered to a 1% false discovery rate (FDR) using the target-decoy strategy combined with linear discriminant analysis. Proteins were quantified only from peptides with a summed Signal to Noise threshold of >=200 and MS2 isolation specificity of 0.5. For each modification, the total summed Signal to Noise was multiplied by the MeCP2 normalization factor for each sample. The log2 fold-change for each treatment condition compared to its control was calculated and displayed in the heatmap.

### Phenotypic characterization of knock-in mice

Consistent with a previous study (27), mice were backcrossed a minimum of 3 generations before undergoing aging characterization and a minimum of 12 generations for behavioral scoring. For aging studies, WT (MeCP2-WT/Y) and QKI (MeCP2-QKI/Y) littermates were housed for a year, noting survival, which was graphed using Kaplan-Meier plots. Mice underwent phenotypic scoring, assessing mobility, gait, hindlimb clasping, tremor, breathing, and general condition, where mice akin to WT are scored 0, 1 with symptoms present, and 2 with severe symptoms as previously described (27). All assessments were performed blinded to genotype.

### MeCP2 chromatin immunoprecipitation sequencing (ChIP-seq)

#### MeCP2 ChIP-seq

MeCP2 ChIP-seq in MeCP2 QKI and WT mice was performed as described in Boxer, Renthal et al., 2020 with minor modifications. Forebrain (cortex + hippocampus) was dissected from 8-to-11-week-old male MeCP2 QKI and WT mice and flash frozen. Tissue was homogenized in 1% formaldehyde and cross-linked by rotating for 10 minutes at room temperature. Cross-linking was quenched with 125 mM glycine for 5 minutes at room temperature. Tissue was pelleted by spinning at 500 x g for 5 minutes at 4°C, washed once with PBS, and spun again at 500 x g for 5 minutes at 4°C. Cells were lysed by resuspending in Buffer L1 (50 mM HEPES pH 7.5, 140 mM NaCl, 1 mM EDTA pH 8, 1 mM EGTA pH 8, 10% glycerol, 0.5% NP-40, 0.25% Triton X-100, 1X protease inhibitors) and rotating for 10 minutes at 4°C. Nuclei were pelleted by spinning at 500 x g for 5 minutes at 4°C, then washed by rotating for 10 minutes at 4°C in Buffer L2 (10 mM Tris pH 8, 200 mM NaCl, 1X protease inhibitors). Nuclei were pelleted by spinning at 500 x g for 5 minutes, then resuspended in Buffer LB3 (10 mM Tris pH 8, 100 mM NaCl, 1 mM EDTA, 0.5 mM EGTA, 0.1% Na-Deoxycholate, 0.5% N-Lauroylsarcosine, 1X protease inhibitors). Chromatin was sonicated in a Diagenode Bioruptor (High power, 65 cycles, 30 seconds on, 45 seconds off). Insoluble material was removed by spinning at 16,000 x g for 10 minutes at 4°C, and Triton X-100 was added to soluble chromatin to a final concentration of 1%. Chromatin was pre-cleared for two hours with Protein A Dynabeads, then incubated with Protein A Dynabeads conjugated to MeCP2 antibody (14) overnight at 4°C. Beads were washed at 4°C twice with Low Salt Buffer (20 mM Tris pH 8, 150 mM NaCl, 2 mM EDTA, 1% Triton X-100, 0.1% SDS), twice with High Salt Buffer (20 mM Tris pH 8, 500 mM NaCl, 2 mM EDTA, 1% Triton X-100, 0.1% SDS), twice with LiCl Wash Buffer (10 mM Tris pH 8, 1 mM EDTA, 1% NP-40, 250 mM LiCl, 1% sodium deoxycholate) and once with TE Buffer (50 mM Tris pH 8, 10 mM EDTA). Chromatin was eluted off beads in TE Buffer with 1% SDS at 65°C for one hour, and crosslinking was reversed overnight at 65°C. Chromatin was treated with RNase A for 30 minutes at 37°C and Proteinase K for 2 hours at 55°C. DNA was phenol-chloroform extracted and purified with the QIAGEN PCR purification kit. MeCP2 ChIP and input libraries were generated using the NuGEN Ovation Ultralow System V2 following manufacturer instructions. Libraries were sequenced on an Illumina Nextseq 500 with 75 bp single-end reads.

#### MeCP2 ChIP-seq analysis

MeCP2 ChIP-seq was analyzed as in Boxer, Renthal et al., 2020 with minor modifications. Sequencing reads were trimmed with Trimmomatic (v0.36) to remove adapters and low-quality sequence (settings: LEADING:5 TRAILING:5 SLIDINGWINDOW:4:20 MINLEN:50). Trimmed reads were mapped to the mm10 genome with Bowtie2 (v2.2.9) with default parameters. PCR duplicate reads were removed with SAMtools (v0.1.19) rmdup. Reads were extended to 250 bp (to approximate fragment length) with UCSC tools (v363) bedextendranges. Reads for each sample were randomly down-sampled to equal numbers using the GNU shuf utility. MeCP2 ChIP-seq and input reads were quantified in mm10 RefSeq gene bodies (excluding the first 3 kb, which is relatively depleted for MeCP2 binding and DNA methylation, and thus excluding genes less than 4.5 kb in length) using Bedtools map (v2.26). For DNA methylation, bisulfite sequencing data from 10-week-old mouse cortex from (51) was used to calculate the DNA methylation density as the number of cytosines/(number of cytosines + thymines) in either the CA or CG dinucleotide context in each gene body. Gene bodies with fewer than 5 total CA, CG, MeCP2 ChIP, or input reads were excluded from analysis. To generate smooth-line correlation plots, genes were sorted by CA or CG DNA methylation density, and a sliding window was defined by 400-gene bins with 40-gene steps. The mean log2 fold-change MeCP2 ChIP/input for each bin was plotted for each MeCP2 QKI or WT sample.

### dLGN Acute Slice preparation for Electrophysiology

Acute sections of brain tissue containing the dLGN as well as the optic tract comprising RGC axons were prepared as previously described (28, 35). Anesthesia of mice was performed through inhalation of isoflurane followed by immediate decapitation. The mouse head was immersed into an ice-cold, oxygenated solution containing (in mM): 130 K-gluconate, 15 KCl, 0.05 EGTA, 20 HEPES, and 25 glucose (pH 7.4 with NaOH; Sigma). A single 250 μm-thick parasagittal slice was cut for each mouse in this ice-cold solution using a sapphire blade (Delaware Diamond Knives, Wilmington, DE) on a vibratome (VT1000S; Leica, Deerfield, IL). Slices were then recovered at 32°C for 20-30 minutes in oxygenated artificial cerebrospinal fluid (ACSF) in mM: 125 NaCl, 26 NaHCO_3_, 1.25 NaH_2_PO_4_, 2.5 KCl, 1.0 MgCl_2_, 2.0 CaCl_2_, and 25 glucose (Sigma), adjusted to 310-312 mOsm with water.

### Electrophysiology

Whole-cell voltage clamp recordings of thalamocortical (TC) neurons in dLGN were performed as previously described (28) with borosilicate glass pipettes (1-2.5 MOhms, Sutter Instrument, Novato, CA) filled with an internal solution (in mM): 35 CsF, 100 CsCl, 10 EGTA, 10 HEPES, and an L-type Ca^2+^ antagonist 0.1 methoxyverapamil (290-300 mOsm, pH 7.3; Sigma). Experiments were performed at room temperature in oxygenated ACSF containing 50 μM of the GABA_A_ receptor antagonist picrotoxin (Tocris, Ellisville, MO). Recordings from synapses that were silent (failure of transmission at -70 mV, but EPSC evident at +40 mV) were averaged for at least 3-5 trials. Single RGC fiber EPSCs were obtained using a threshold method as previously described (13, 24).

For Paired Pulse Ratio (PPR) experiments, the optic tract was stimulated twice with randomized inter-stimulus intervals (ISIs) of 50, 100, 250, and 500 ms while holding the cell at -70 mV. PPR was calculated by dividing A2/A1, where A2 is the peak amplitude of the second EPSC and A1 is the peak amplitude of the first EPSC. For ISIs of 50, 100, and 250 ms, the average waveform of the first EPSC at an ISI of 500 ms was subtracted from the second EPSC to obtain a more accurate measure of A2. Averages of three to six trials of each ISI were used to calculate PPR. To inhibit modulatory and postsynaptic contributions to short-term synaptic plasticity at the retinogeniculate synapse (52–54), recordings were performed in the presence of (in μM): 5 CGP-55845 (GABA_B_-receptor antagonist), 10 cyanopindolol (5-HT_1_ receptor antagonist), 50 cyclothiazide (AMPAR desensitization inhibitor), 10 DPCPX (A1 adenosine receptor antagonist), and 50 LY-341495 (mGluR antagonist).

### Fiber Fraction

The fiber fraction (FF = single fiber current amplitude ÷ maximal current amplitude) was calculated to approximate RGC convergence onto each TC neuron in the dLGN. Maximal current amplitudes (“Max”) were determined by stimulating the optic tract at increasing intensities ≥50 μA until the amplitude of the EPSC reached a plateau. Prior to recording EPSCs for each cell, a stimulating electrode filled with ACSF was moved along the optic tract in order to determine a site that would activate as many RGC axons as possible and yield the largest maximal EPSC current. A second electrode was positioned just above the surface of the slice to serve as a local ground. The stimulus intensity was then decreased systematically until a failure of synaptic transmission was observed, then increased by 0.25 μA until an EPSC was visible whose amplitude is the single fiber amplitude (“SF”). The fiber fraction is the SF amplitude divided by the maximal amplitude (FF = SF ÷ Max). For each stimulus intensity, we recorded the synaptic response at holding potentials of both -70 mV and +40 mV with an inter-trial interval of 20 seconds, yielding at least two fiber fraction values for each cell recorded. To determine NMDAR-mediated amplitudes, the more slowly rising peak NMDAR-mediated EPSCs were measured following the initial decay of the AMPAR transient. In some cases, only the minimal threshold response was quantified for SF amplitude because it was difficult to distinguish between the recruitment of an additional fiber from trial-to-trial variation of the same fiber.

However, if an incremental step in stimulation intensity of 0.25 μA recruited an EPSC with at least five times the amplitude of the initial SF EPSC at either -70 mV or +40 mV, this second SF amplitude was included in our analysis (“SF_2_”). Our reasoning for this 5x cutoff is the inability to confidently attribute a modestly increased EPSC to the activation of a second RGC axon; we cannot exclude the possibility of stochastic variations in vesicular release of the initial SF. The 5x cutoff was determined after subtracting the first SF amplitude from the second SF amplitude. For “silent synapses”, SF_2_ was always included in our analysis regardless of amplitude because SF_1_ at -70mV = 0. Additional details regarding the justification of fiber fractions are available in the supplementary sections of Hooks and Chen, 2008 and Noutel et al., 2011.

### Western Blotting

Tissue for Western blotting was dissected as described above for electrophysiology, in an ice-cold potassium gluconate solution. Following anesthesia and decapitation, the most anterior portion of the brain containing the olfactory bulb was cut and discarded using a razor blade, creating a flat surface for the remainder of the brain that was superglued onto a Leica VT1000s vibratome. Three 250-300 μm coronal sections of the most posterior visual cortex were cut and gently placed into a 24-well plate with ice cold potassium gluconate. Sections continued to be collected, from posterior to anterior, until sections containing dLGN were visible. The dLGN, as well as the most dorsal portions of visual cortex, were microdissected out with a scalpel under a dissecting microscope, placed into separate tubes, and flash frozen with liquid nitrogen.

Flash-frozen tissue was homogenized in ice-cold RIPA buffer (50 mM Tris-HCl pH 8.0, 150 mM NaCl, 1% NP-40, 0.5% sodium deoxycholate, 0.1% sodium dodecyl sulfate, protease inhibitors, and 1% Phosphatase Inhibitor Cocktail 2 and 3 (Sigma)) and incubated on ice for 15 minutes. Sonication of lysates was performed using a Diagenode Bioruptor on high power for 10 cycles (30 seconds on then 30 seconds off), followed by centrifugation (16,000 x g, 10 minutes, 4°C) to pellet the insoluble fraction. The supernatant was carefully transferred to a new tube, then Nupage LDS 4x sample buffer was added to a final concentration of 1x, as well as 2-mercaptoethanol (5% final concentration). Samples were boiled at 95°C for fifteen minutes, electrophoresed on 4-12% Bis-tris gels (Life Technologies), and transferred to nitrocellulose membranes, which were blocked with 5% dry milk in TBS-T for one hour at room temperature. Membranes were incubated with primary antibodies overnight at 4°C, washed three times with TBS-T, incubated with IR dye secondary antibodies for one hour, and again washed with TBS-T three times. Western blots were imaged using the Li-Cor Odyssey platform and quantified using ImageJ/Fiji. The following primary antibodies and dilutions were used: custom anti-pS421 MeCP2 antibody (1:1000, rabbit, generated in the Greenberg lab in Zhou et al., 2006), anti-total MeCP2 N-terminal antibody (1:2000, mouse, clone Men-8 from Sigma-Aldrich), and ß-actin (1:2000, mouse, Abcam ab8226).

### Late dark rearing for bulk RNA sequencing

Mice were housed under standard conditions in a room set to a 12-hour light/dark cycle, with food and water *ad libitum*. Late dark rearing (LDR) experiments were performed as previously described (24, 35), with cages of mice reared in the aforementioned conditions until P20, then placed in a custom-designed, ventilated, light-proof cabinet located in a dark room using night vision goggles (Pulsar). Mice were then re-exposed to light at P27, by moving their cages to a brightly lit separate chamber in the cabinet for four uninterrupted hours. Unstimulated control mice remained in the dark chamber and were euthanized by isoflurane and decapitated in a dark room to limit the induction of experience-dependent gene expression upon exposure to light. Lights were turned on immediately after the decapitated head was submerged in an ice-cold potassium gluconate solution. An acute slice preparation as described above was performed to dissect out the dLGN and visual cortex for RT-qPCR or bulk RNA-sequencing.

### RNA isolation for RT-qPCR

Tissue samples were triturated with a needle and syringe with TRIzol (Life Technologies, 15596026) to subsequently harvest RNA. The RNeasy Micro Kit (Qiagen, 74004) was then used to purify RNA. RNA (120 ng) across all samples were converted to cDNA using the High-Capacity cDNA Kit (Thermo Fisher, 4368813) according to manufacturer’s instructions. cDNA was diluted by 20-fold and standard RT-qPCR was performed using SYBR Green Master Mix (Thermo Fisher, A25743). qPCR was performed with technical triplicates using a QuantStudio 3 qPCR machine (Applied Biosystems). qPCR target gene expression was normalized against the housekeeping gene *Gapdh*. Expression for each qPCR target gene was normalized to the housekeeping gene *Gapdh*. The following sets of primers were used to amplify the genes of interest.

Primer sets:

*Gapdh* F: AGGTCGGTGTGAACGGATTTG

*Gapdh* R: GGGGTCGTTGATGGCAACA

*Npas4* F: ACCTAGCCCTACTGGACGTT

*Npas4* R: CGGGGTGTAGCAGTCCATAC

*Fn14* F: GACCTCGACAAGTGCATGGACT

*Fn14* R: CGCCAAAACCAGGACCAGACTA

*Fos* F: GGCAGAAGGGGCAAAGTAGA

*Fos* R: GCTGCAGCCATCTTATTCCG

*Bdnf* F: GATGCCGCAAACATGTCTATGA

*Bdnf* R: AATACTGTCACACACGCTCAGCTC

### Single-nucleus RNA-seq (SnRNA-seq)

#### SnRNA-seq

To isolate dLGN nuclei, we combined flash-frozen dLGN tissue dissected from P20 mice in artificial cerebrospinal fluid (ACSF) (in mM): 125 NaCl, 26 NaHCO_3_, 1.25 NaH_2_PO_4_, 2.5 KCl, 1.0 MgCl_2_, 2.0 CaCl_2_, and 25 glucose. Three male mice per genotype (MeCP2-WT/Y and MeCP2-QKI/Y) were dissected for each replicate, for a total of 12 mice dissected over two replicates. To the tissue, we added 1 mL of buffer HB (0.25 M sucrose, 25 mM KCl, 5 mM MgCl_2_, 20 mM Tricine-KOH, pH 7.8, supplemented with RNase inhibitors, 1 mM DTT, 0.15 mM spermine, 0.5 mM spermidine) and homogenized with a Dounce 5X with a loose pestle and 10X with a tight pestle. 5% IGEPAL CA-630 (32 μL) was added prior to homogenization with a tight pestle 5 more times and filtering through a 40-μm strainer into a 50 mL conical collection tube. 1 mL working solution (50% iodixanol, 25 mM KCl, 5 mM MgCl_2_, 20 mM Tricine-KOH, pH 7.8, supplemented with RNase inhibitors, DTT, spermine and spermidine) were added. Homogenized tissue was gently layered on top of 100 μL of 30% iodixanol on top of a layer of 100 μL of 40% iodixanol (diluted from working solution). Samples were centrifuged at 1,500 x g for 15min. Nuclei (50 μL) were collected from the 30/40% iodixanol interface. An aliquot of each sample was incubated with trypan blue, and nuclei were counted using a standard hemocytomter, for a total of ∼10,000 nuclei per pooled sample.

SnRNA-seq was performed using the 10X Genomics Chromium Next GEM Single Cell 3’ v3.1 (Dual Index) kit. Each reaction lane was loaded with up to 10,000 nuclei from each pooled dLGN sample. Subsequent steps for cDNA amplification and library preparation were conducted according to the manufacturer’s protocol (10X Genomics). Samples were sequenced using Illumina NextSeq500 with 28 bp (R1), 90 bp (R2), and 10 bp (index). Our final dataset includes 34,693 total nuclei collected from WT and QKI samples.

#### SnRNA-seq analysis

FASTQ files were created using the standard bcl2fastq pipeline from Illumina. Gene expression tables for each nuclear barcode were generated via the CellRanger 3.0.0 pipeline as designed by 10X Genomics. Samples were demultiplexed, and all QKI or WT samples were merged using the CellRanger aggr function using default parameters. The datasets were loaded into R and analyzed using the Seurat (v3) package. Nuclei were removed from the dataset if they contained fewer than 500 detected genes, displayed more than 5% of reads mapping to mitochondrial genes, or had RNA counts detected at a level greater than 2 standard deviations higher than the mean value in their assigned cell type (likely reflecting doublets and multiplets).

All nuclei in either QKI or WT samples were considered together for clustering and dimensionality reduction. Data was normalized using Seurat’s (v3) NormalizeData, and the 2,000 top variable genes across nuclei were identified using FindVariableFeatures function. Data was then integrated using FindIntegrationAnchors (16 dimensions) and IntegrateData (16 dimensions). Data was then scaled using ScaleData, and principal component analysis using the RunPCA function (npcs = 16) was performed. A shared nearest neighbor graph was constructed using the FindNeighbors function (considering the top 16 principal components), and clustering was assigned using the FindClusters function (resolution = 0.5). After, clusters with fewer than 100 nuclei were removed.

The following marker genes were used to assign cell type clusters identified: pan-neuronal (*Rbfox3*); pan-excitatory neurons (*Slc17a7*); pan-inhibitory neurons (*Gad1*); oligodendrocytes (*Mag*); oligodendrocyte precursor cells (*Pdgfra*); astrocytes (*Aldoc*); microglia (*Cx3cr1*); endothelial cells (*Cldn5*).

Differential gene expression analysis between QKI and WT cells within each cluster was conducted using Wilcoxon Rank Sum test via the FindMarkers function (default parameters), and significant genes were defined as those with an adjusted p-value less than 0.05.

### RNA-Sequencing (RNA-seq)

#### RNA-seq

Tissue was dissected from WT and QKI male littermates (MeCP2-WT/Y and MeCP2-QKI/Y) in either ACSF (P16, P20 datasets) or the solution used for electrophysiology experiments (P15, P30, and LDR datasets). Tissue was homogenized in Trizol reagent, and RNA was chloroform extracted and then purified using the Qiagen RNeasy Micro Kit with on-column DNase treatment according to the manufacturer’s instructions. Total RNA (150-200 ng for dLGN libraries, 500 ng for visual cortex libraries) was used to generate libraries following rRNA depletion (NEBNext, E6310X) according to the manufacturer’s instructions (NEBNext, E7420). Libraries were sequenced on an Illumina Nextseq 500 with 85 bp single-end reads. All libraries that were compared as a set for differential gene expression analysis were prepared and sequenced at the same time to reduce batch effects, before being subsequently analyzed with our standardized RNA-seq data analysis pipeline (below).

#### RNA-seq analysis

Analysis was performed as described previously (49). Briefly, reads were trimmed with Trimmomatic (v0.36) (55) to remove Illumina adaptors and low-quality sequence (settings: LEADING:5 TRAILING:5 SLIDINGWINDOW:4:20 MINLEN:50). Trimmed reads were mapped to the mm10 Refseq transcriptome and genome using STAR (v2.5.2b) (56). Reads were counted in the full gene body (TSS to TTS of longest isoform, in order to include intronic reads) using Subread featureCounts. Differential expression was performed with the R package edgeR (v3.34.1) (57) or DESeq2 (v1.34.0) (58). Genes with low counts were filtered by keeping only genes with rowSums(cpm(y)>1) >= (number of samples per group). Differentially expressed genes were defined by an FDR < 0.05 with no fold-change cut-off. For DESeq2 analyses, shrinkage of effect sizes was performed using lfcshrink and the apeglm method following differential expression analysis (59).

### Eye-specific segregation in the LGN

QKI mice and littermate WT controls (P58-64) were anesthetized with isoflurane and injected intra-ocularly with 2 µL of a 2% solution of cholera toxin subunit B conjugated to Alexa Fluor 488 or Alexa Fluor 555. Four days after injection, mice were anesthetized with ketamine/xylazine and transcardially perfused with 4% paraformaldehyde. Coronal sections (75 µm) were mounted with mounting medium containing DAPI. Non-saturated 10x images were acquired with an Olympus DP72 camera on an Olympus BX63 fluorescent microscope. Images were analyzed using Fiji using a previously described threshold-independent method (13, 37). Background signal was subtracted and the dorsal LGN was selected. The logarithm of the ratio of ipsilateral to contralateral channel intensity (R) was calculated for each pixel, and then the mean variance of R was calculated for each section. The mean variance of R was used to compare eye-specific segregation across genotype.

## ACKNOWLEDGEMENTS

We thank Drs. Hume Stroud, Elizabeth Pollina, Lucas Cheadle, Christopher Davis, David Harmin, Thomas Schwarz, Bruce Bean, Pascal Kaeser, David Ginty, Gord Fishell, Lisa Goodrich, and all members of the Greenberg and Chen labs for discussions and input on the project. This work was supported by the Rett Syndrome Research Trust, NIH R01NS048276 (M.E.G.), R01EY013613 (C.C.), K99NS112415 (L.D.B), the Stuart H.Q. & Victoria Quan Fellowship (T.W.), and the Harvard Department of Neurobiology Graduate Fellowship (T.W.).

## Author Contributions

Experiments were designed by C.P.T., T.W., L.D.B, W.R., S.T., C.C., and M.E.G. L.D.B., W.R., E.L., A.S., and R.S. generated the quadruple knock-in mouse, did initial characterization of QKI mice, and performed mass spectrometry experiments. C.P.T. performed all electrophysiology experiments. C.P.T. (P30 & LDR), T.W. (P15, P16, & P20), and K.M. performed all gene expression experiments. S.T. performed eye-specific segregation experiments. C.L. performed RT-qPCR validation of the LDR and light exposure paradigm. The manuscript was written by C.P.T., T.W., L.D.B., E.C.G., C.C., and M.E.G.

## REFERENCES

1. R. E. Amir, et al., Rett syndrome is caused by mutations in X-linked MECP2, encoding methyl-CpG-binding protein 2. Nat. Genet. 23, 185–188 (1999).

2. J. D. Lewis, et al., Purification, sequence, and cellular localization of a novel chromosomal protein that binds to Methylated DNA. Cell 69, 905–914 (1992).

3. M. Chahrour, H. Y. Zoghbi, The Story of Rett Syndrome: From Clinic to Neurobiology. Neuron 56, 422–437 (2007).

4. H. T. Chao, H. Y. Zoghbi, C. Rosenmund, MeCP2 Controls Excitatory Synaptic Strength by Regulating Glutamatergic Synapse Number. Neuron 56, 58–65 (2007).

5. Y. Asaka, D. G. M. Jugloff, L. Zhang, J. H. Eubanks, R. M. Fitzsimonds, Hippocampal synaptic plasticity is impaired in the Mecp2-null mouse model of Rett syndrome. Neurobiol. Dis. 21, 217–227 (2006).

6. E. S. Na, et al., A mouse model for MeCP2 duplication syndrome: MeCP2 overexpression impairs learning and memory and synaptic transmission. J. Neurosci. 32, 3109–3117 (2012).

7. Z. Qiu, et al., The Rett syndrome protein MeCP2 regulates synaptic scaling. J. Neurosci. 32, 989–994 (2012).

8. M. P. Blackman, B. Djukic, S. B. Nelson, G. G. Turrigiano, A critical and cell-autonomous role for MeCP2 in synaptic scaling up. J. Neurosci. 32, 13529–13536 (2012).

9. M. C. Crair, Neuronal activity during development: Permissive or instructive? Curr. Opin. Neurobiol. 9, 88–93 (1999).

10. A. D. Huberman, M. B. Feller, B. Chapman, Mechanisms Underlying Development of Visual Maps and Receptive Fields. Annu. Rev. Neurosci. 31, 479–509 (2008).

11. N. Ballas, D. T. Lioy, C. Grunseich, G. Mandel, Non-cell autonomous influence of MeCP2-deficient glia on neuronal dendritic morphology. Nat. Neurosci. 12, 311–317 (2009).

12. C. A. Chapleau, et al., Dendritic spine pathologies in hippocampal pyramidal neurons from Rett syndrome brain and after expression of Rett-associated MECP2 mutations. Neurobiol. Dis. 35, 219–233 (2009).

13. J. Noutel, Y. K. Hong, B. Leu, E. Kang, C. Chen, Experience-Dependent Retinogeniculate Synapse Remodeling Is Abnormal in MeCP2-Deficient Mice. Neuron 70, 35–42 (2011).

14. W. G. Chen, et al., Derepression of BDNF Transcription Involves Calcium-Dependent Phosphorylation of MeCP2. Science (80-.). 302, 885–889 (2003).

15. D. Damen, R. Heumann, MeCP2 phosphorylation in the brain: From transcription to behavior. Biol. Chem. 394, 1595–1605 (2013).

16. E. Bellini, et al., MeCP2 post-translational modifications: A mechanism to control its involvement in synaptic plasticity and homeostasis? Front. Cell. Neurosci. 8, 1–15 (2014).

17. L. Chen, et al., MeCP2 binds to non-CG methylated DNA as neurons mature, influencing transcription and the timing of onset for Rett syndrome. Proc. Natl. Acad. Sci. U. S. A. 112, E2982 (2015).

18. D. H. Ebert, et al., Activity-dependent phosphorylation of MeCP2 threonine 308 regulates interaction with NCoR. Nature 499, 341–345 (2013).

19. M. L. Gonzales, S. Adams, K. W. Dunaway, J. M. LaSalle, Phosphorylation of Distinct Sites in MeCP2 Modifies Cofactor Associations and the Dynamics of Transcriptional Regulation. Mol. Cell. Biol. 32, 2894–2903 (2012).

20. G. Stefanelli, et al., Brain phosphorylation of MeCP2 at serine 164 is developmentally regulated and globally alters its chromatin association. Sci. Rep. 6, 1–15 (2016).

21. J. Tao, et al., Phosphorylation of MeCP2 at serine 80 regulates its chromatin association and neurological function. Proc. Natl. Acad. Sci. U. S. A. 106, 4882–4887 (2009).

22. Z. Zhou, et al., Brain-Specific Phosphorylation of MeCP2 Regulates Activity-Dependent Bdnf Transcription, Dendritic Growth, and Spine Maturation. Neuron 52, 255–269 (2006).

23. S. Cohen, et al., Genome-Wide Activity-Dependent MeCP2 Phosphorylation Regulates Nervous System Development and Function. Neuron 72, 72–85 (2011).

24. B. M. Hooks, C. Chen, Distinct Roles for Spontaneous and Visual Activity in Remodeling of the Retinogeniculate Synapse. Neuron 52, 281–291 (2006).

25. B. M. Hooks, C. Chen, Vision triggers an experience-dependent sensitive period at the retinogeniculate synapse. J. Neurosci. 28, 4807–4817 (2008).

26. J. Guy, H. Cheval, J. Selfridge, A. Bird, The Role of MeCP2 in the Brain. Annu. Rev. Cell Dev. Biol. 27, 631–652 (2011).

27. J. Guy, J. Gan, J. Selfridge, S. Cobb, A. Bird, Reversal of Neurological Defects in a Mouse Model of Rett Syndrome. Science (80-.). 315, 1143–1147 (2007).

28. C. Chen, W. G. Regehr, Developmental remodeling of the retinogeniculate synapse. Neuron 28, 955–966 (2000).

29. E. Y. Litvina, C. Chen, Functional Convergence at the Retinogeniculate Synapse. Neuron 96, 330–338.e5 (2017).

30. Y. K. Hong, C. Chen, Wiring and rewiring of the retinogeniculate synapse. Curr. Opin. Neurobiol. 21, 228–237 (2011).

31. X. Liu, C. Chen, Different roles for AMPA and NMDA receptors in transmission at the immature retinogeniculate synapse. J. Neurophysiol. 99, 629–643 (2008).

32. J. Demas, S. J. Eglen, R. O. L. Wong, Developmental loss of synchronous spontaneous activity in the mouse retina is independent of visual experience. J. Neurosci. 23, 2851–2860 (2003).

33. M. B. Feller, D. P. Wellis, D. Stellwagen, F. S. Werblin, C. J. Shatz, Requirement for cholinergic synaptic transmission in the propagation of spontaneous retinal waves. Science (80-.). 272, 1182–1187 (1996).

34. M. Meister, R. L. Wong, D. a Baylor, C. J. Shatz, Synchronous bursts of action potentials in ganglion cells of the developing mammalian retina. Science (80-.). 252, 939–943 (1991).

35. L. Jaubert-Miazza, et al., Structural and functional composition of the developing retinogeniculate pathway in the mouse. Vis. Neurosci. 22, 661–676 (2005).

36. C. L. Torborg, M. B. Feller, Unbiased analysis of bulk axonal segregation patterns. J. Neurosci. Methods 135, 17–26 (2004).

37. M. E. Greenberg, E. B. Ziff, Stimulation of 3T3 cells induces transcription of the c-fos proto-oncogene. Nature 311, 433–438 (1984).

38. E. L. Yap, et al., Bidirectional perisomatic inhibitory plasticity of a Fos neuronal network. Nature 590, 115–121 (2021).

39. Y. Lin, et al., Activity-dependent regulation of inhibitory synapse development by Npas4. Nature 455, 1198–1204 (2008).

40. B. L. Bloodgood, N. Sharma, H. A. Browne, A. Z. Trepman, M. E. Greenberg, The activity-dependent transcription factor NPAS4 regulates domain-specific inhibition. Nature 503, 121–125 (2013).

41. I. Spiegel, et al., Npas4 regulates excitatory-inhibitory balance within neural circuits through cell-type-specific gene programs. Cell 157, 1216–1229 (2014).

42. P. B. Shieh, S. C. Hu, K. Bobb, T. Timmusk, A. Ghosh, Identification of a signaling pathway involved in calcium regulation of BDNF expression. Neuron 20, 727–740 (1998).

43. X. Tao, S. Finkbeiner, D. B. Arnold, A. J. Shaywitz, M. E. Greenberg, Ca2+ influx regulates BDNF transcription by a CREB family transcription factor-dependent mechanism. Neuron 20, 709–726 (1998).

44. E. J. Hong, A. E. McCord, M. E. Greenberg, A Biological Function for the Neuronal Activity-Dependent Component of Bdnf Transcription in the Development of Cortical Inhibition. Neuron 60, 610–624 (2008).

45. L. Cheadle, et al., Visual Experience-Dependent Expression of Fn14 Is Required for Retinogeniculate Refinement. Neuron 99, 525–539.e10 (2018).

46. T. Stuart, et al., Comprehensive Integration of Single-Cell Data. Cell 177, 1888–1902.e21 (2019).

47. B. T. Kalish, et al., Single-cell transcriptomics of the developing lateral geniculate nucleus reveals insights into circuit assembly and refinement. Proc. Natl. Acad. Sci. U. S. A. 115, E1051–E1060 (2018).

48. K. Krishnan, et al., MeCP2 regulates the timing of critical period plasticity that shapes functional connectivity in primary visual cortex. Proc. Natl. Acad. Sci. U. S. A. 112, E4782–E4791 (2015).

49. L. D. Boxer, et al., MeCP2 Represses the Rate of Transcriptional Initiation of Highly Methylated Long Genes. Mol. Cell 77, 294–309 (2020).

50. N. Picard, M. Fagiolini, MeCP2: an epigenetic regulator of critical periods. Curr. Opin. Neurobiol. 59, 95–101 (2019).

51. J. Guy, B. Hendrich, M. Holmes, J. E. Martin, A. Bird, A mouse Mecp2-null mutation causes neurological symptoms that mimic rett syndrome. Nat. Genet. 27, 322–326 (2001).

52. R. Lister, et al., Global epigenomic reconfiguration during mammalian brain development. Science (80-.). 341, 1237905–1-1237905–12 (2013).

53. C. Chen, D. M. Blitz, W. G. Regehr, Contributions of receptor desensitization and saturation to plasticity at the retinogeniculate synapse. Neuron 33, 779–788 (2002).

54. J. L. Hauser, X. Liu, E. Y. Litvina, C. Chen, Prolonged synaptic currents increase relay neuron firing at the developing retinogeniculate synapse. J. Neurophysiol. 112, 1714–1728 (2014).

55. J. D. S. Reggiani, et al., Brainstem serotonin neurons selectively gate retinal information flow to thalamus 1 2. Neuron, 1–16 (2023).

56. A. M. Bolger, M. Lohse, B. Usadel, Trimmomatic: A flexible trimmer for Illumina sequence data. Bioinformatics 30, 2114–2120 (2014).

57. A. Dobin, et al., STAR: Ultrafast universal RNA-seq aligner. Bioinformatics 29, 15–21 (2013).

58. M. D. Robinson, D. J. McCarthy, G. K. Smyth, edgeR: A Bioconductor package for differential expression analysis of digital gene expression data. Bioinformatics 26, 139–140 (2009).

59. M. I. Love, W. Huber, S. Anders, Moderated estimation of fold change and dispersion for RNA-seq data with DESeq2. Genome Biol. 15, 1–21 (2014).

60. A. Zhu, J. G. Ibrahim, M. I. Love, Heavy-Tailed prior distributions for sequence count data: Removing the noise and preserving large differences. Bioinformatics 35, 2084–2092 (2019).

