## Supporting Information Figures S1 to S5 for "Activity-Induced MeCP2 Phosphorylation Regulates Retinogeniculate Synapse Refinement"

**This PDF file includes:**

Figures S1 to S5

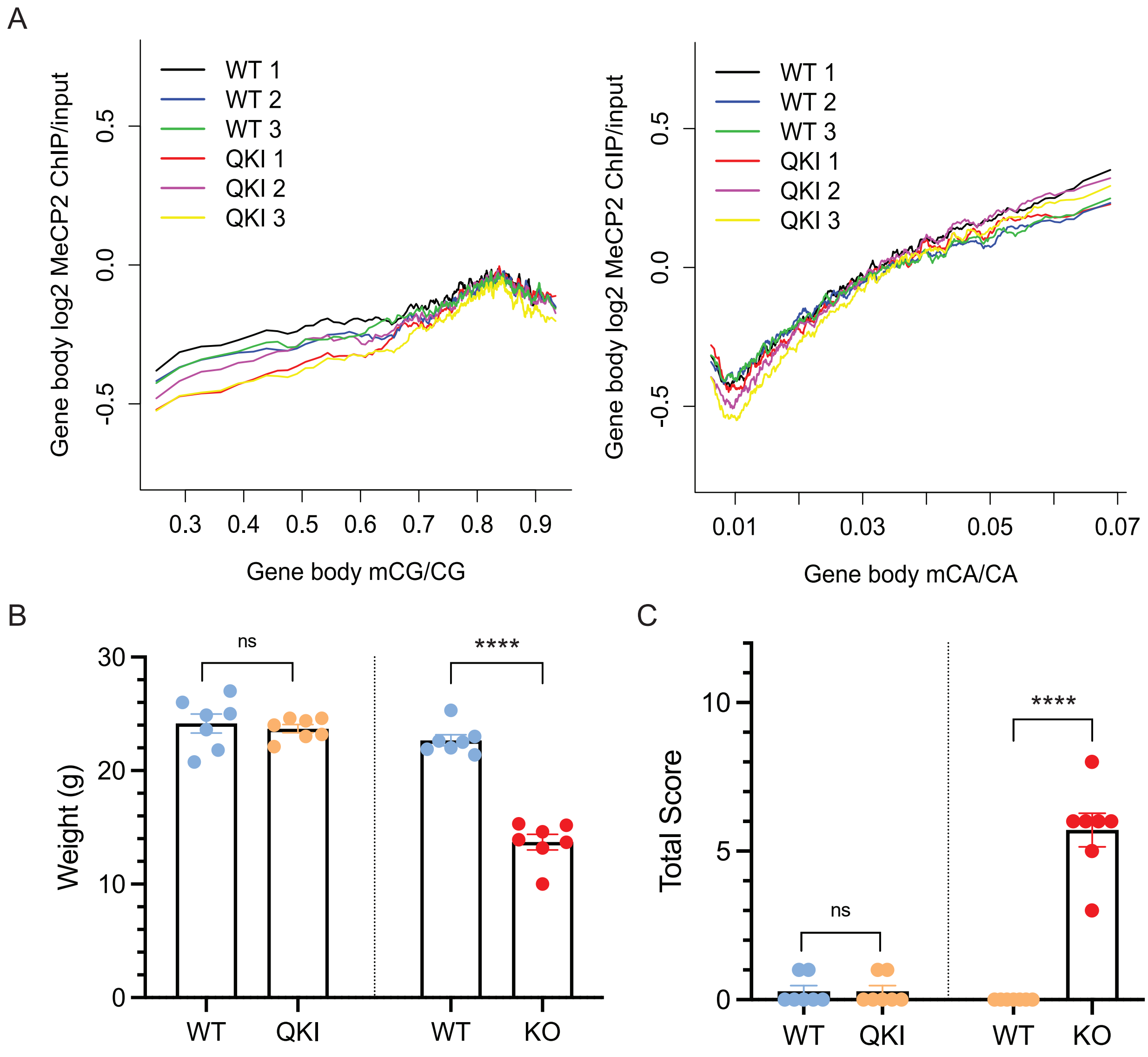


**Figure S1. Validation of quadruple knock-in (QKI) mice.**

**A)** MeCP2 exhibits similar binding to methylated DNA in QKI and WT mice. mCG (*Left*) and mCA (*Right*).

**B)** QKI mice show no differences in weight as compared to WT littermates (n=7). Male littermates were weighed between 6 and 9 weeks of age. Statistics were performed with an unpaired, two-sided, parametric t-test. ns, $p > 0.05$.

**C)** QKI mice show no differences in Rett Syndrome-like phenotypes, as assessed by the Bird scoring method. Male littermates were assessed between the ages of 6 and 9 weeks of age (n=7). Statistics were performed with an unpaired, one-sided, parametric t-test. ns, $p > 0.05$.

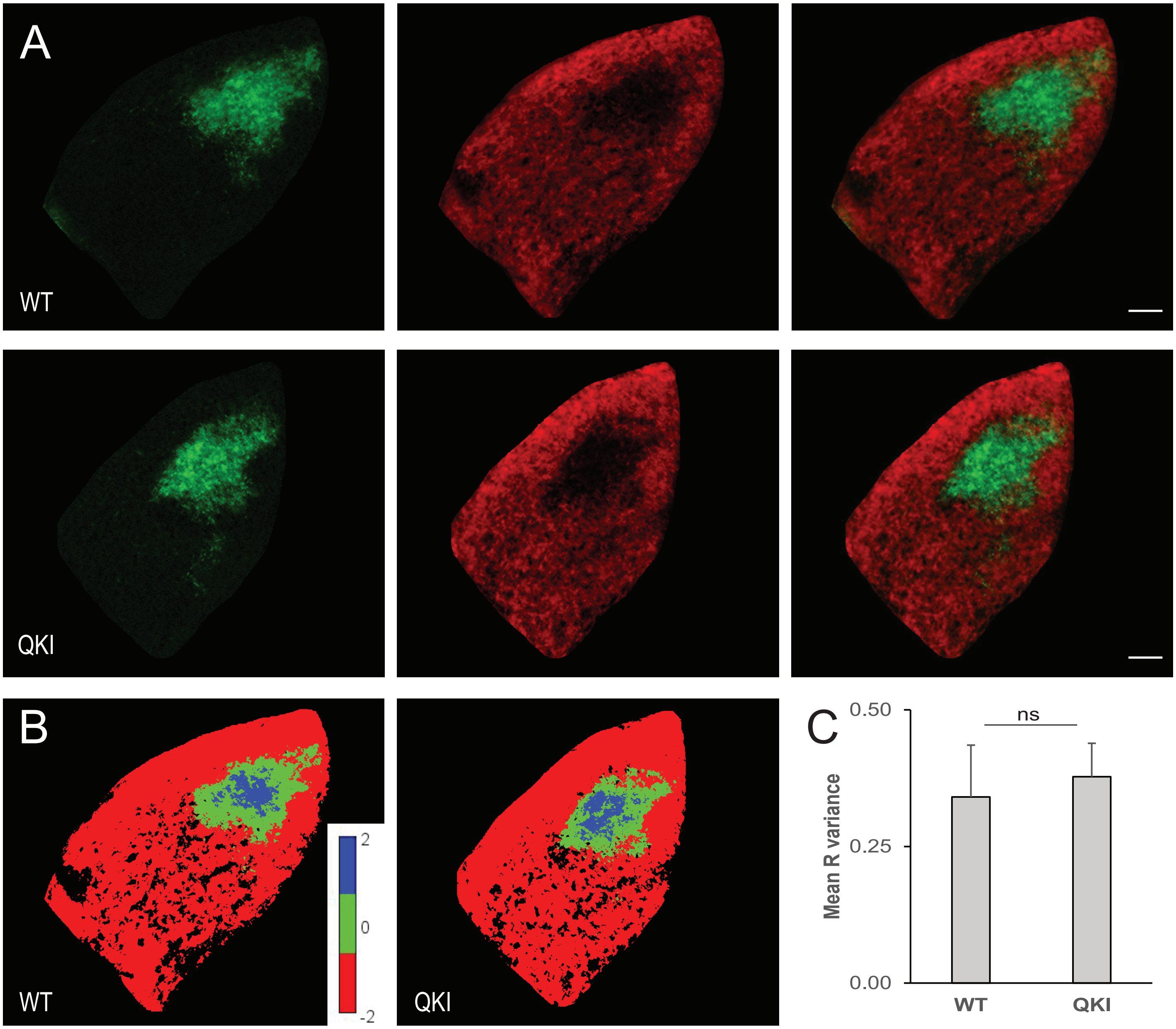


**Figure S2. Eye-specific segregation is normal in QKI adult animals.**
**A)** Representative images of coronal sections of the LGN from P58-P64 QKI and littermate WT control mice with labeled retinal projections from the ipsilateral eye (green) and contralateral eye (red).
**B)** Representative images pseudocolored based on R-values, in which ipsilateral-dominated pixels are blue and contralateral-dominated pixels are red.
**C)** Mean R-variance is not significantly different between WT and QKI mice (student t-test, $p=0.20$). $n=16$ sections from 4 animals for each genotype. Scale bar = 100 µm. Error bars = standard deviation.

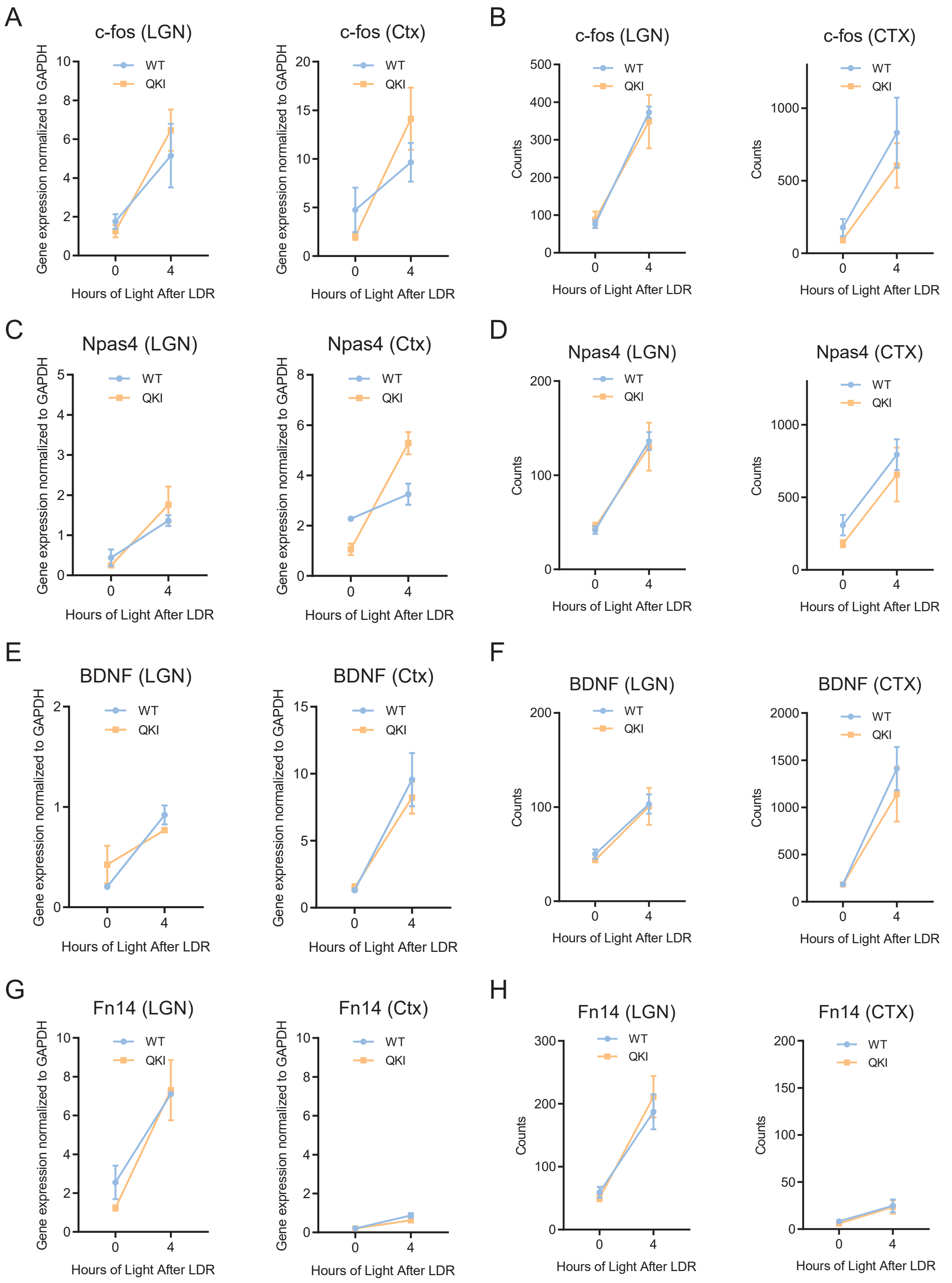


**Figure S3**. **Validation of late dark rearing (LDR) and light exposure paradigm.**
**A-B)** Induction of Fos in dLGN (*Left*) and visual cortex (*Right*) following LDR and 4 hours of light exposure by RT-qPCR (*A*) and by bulk RNA-Seq (*B*).
**C-D)** Induction of Npas4 in dLGN (*Left*) and visual cortex (*Right*) following LDR and 4 hours of light exposure by RT-qPCR (*C*) and by bulk RNA-Seq (*D*).
**E-F)** Induction of Bdnf in dLGN (*Left*) and visual cortex (*Right*) following LDR and 4 hours of light exposure by RT-qPCR (*E*) and by bulk RNA-Seq (*F*).
**G-H)** Induction of Tnfrsf12a (Fn14) in dLGN (*Left*) and visual cortex (*Right*) following LDR and 4 hours of light exposure by RT-qPCR (*G*) and by bulk RNA-Seq (*H*).

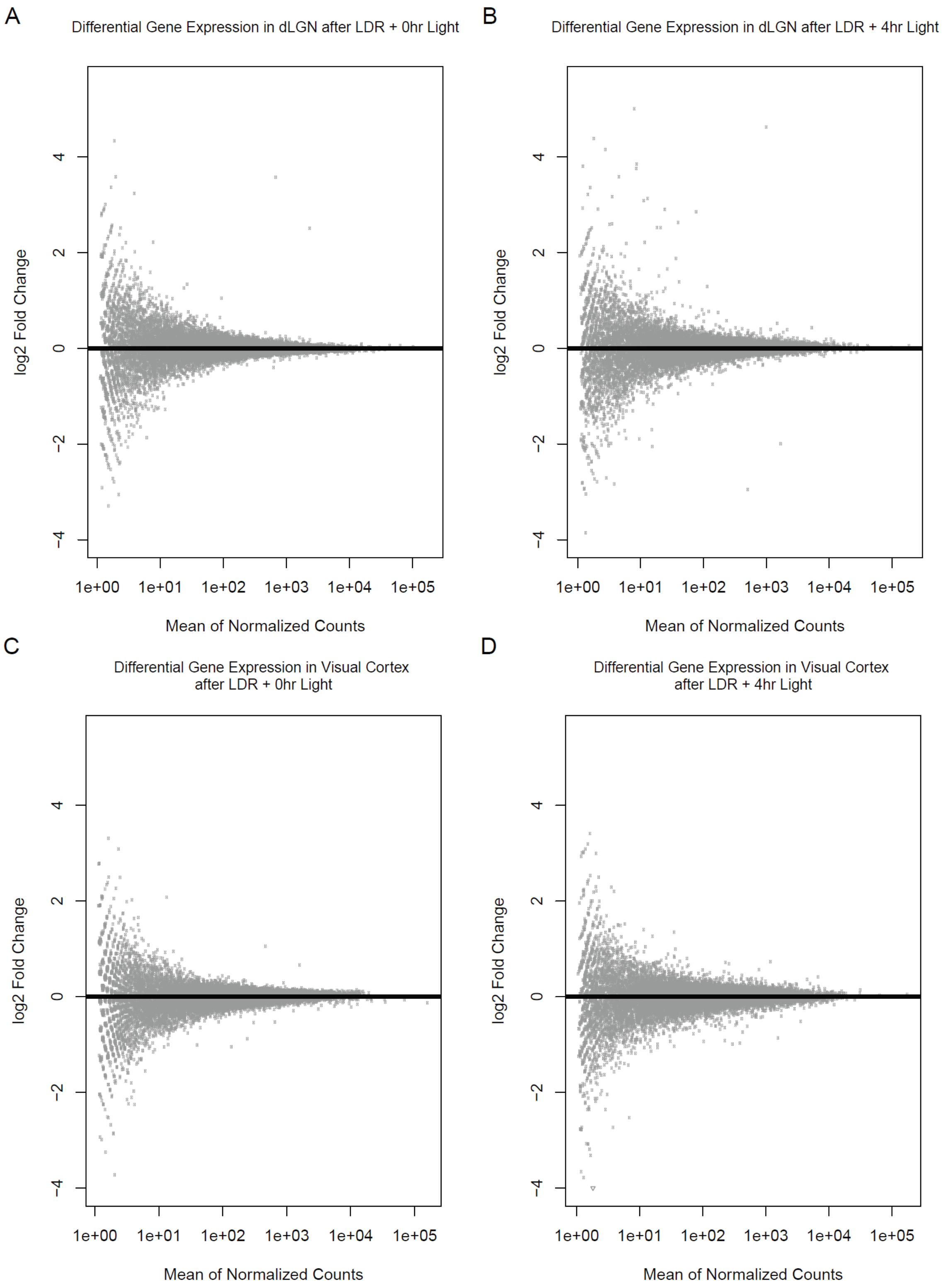


**Figure S4. Late dark rearing (LDR) and light exposure did not reveal detectable activity-induced gene expression differences between wild-type and MeCP2 QKI mice.**

A) Comparison of mean normalized counts versus $log_2$ fold-changes for all genes between WT and QKI dLGN from mice after undergoing LDR and no light exposure reveals no significantly differentially expressed genes.

B) Comparison of mean normalized counts versus $log_2$ fold-changes for all genes between WT and QKI dLGN from mice after undergoing LDR and 4 hours of light exposure reveals no significantly differentially expressed genes.

C) Comparison of mean normalized counts versus $log_2$ fold-changes for all genes between WT and QKI visual cortices from mice after undergoing LDR and no light exposure reveals no significantly differentially expressed genes.

D) Comparison of mean normalized counts versus $log_2$ fold-changes for all genes between WT and QKI visual cortices from mice after undergoing LDR and 4 hours of light exposure reveals no significantly differentially expressed genes.

n=4 for all conditions. $p > 0.05$, Wald test.

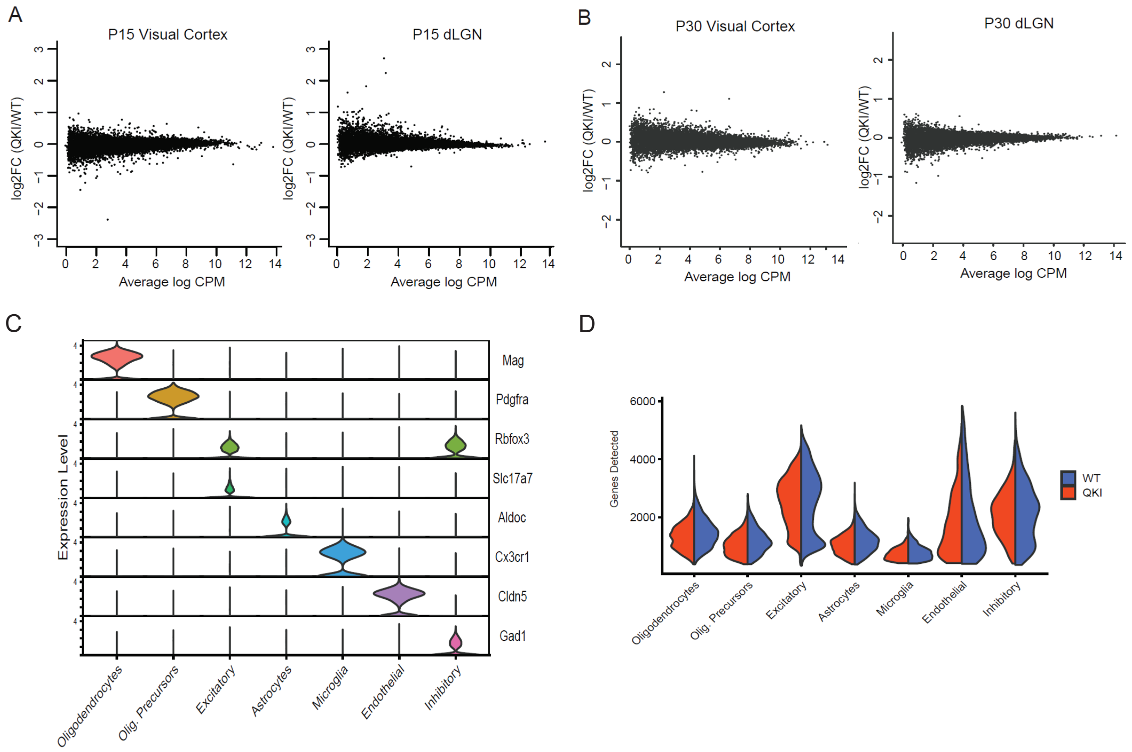


**Figure S5. Gene expression analysis in QKI mice.**

**A)** Mean-difference plot of differential expression analysis from bulk RNA-sequencing of 6 WT and 6 QKI visual cortex (*Left*) and dLGN (*Right*) at P15. No significant differentially expressed genes were detected. Differentially expressed genes were defined by an FDR < 0.05 with no fold-change cut-off. Fisher's exact test with Benjamini-Hochberg correction.

**B)** Mean-difference plot of differential expression analysis from bulk RNA-sequencing of 6 WT and 6 QKI visual cortex (*Left*) and dLGN (*Right*) at P30. No significant differentially expressed genes were detected. p >0.05, Fisher's exact test with Benjamini-Hochberg correction.

**C)** Cell type assignment in P20 dLGN snRNA-seq dataset using the indicated marker genes. The y-axis indicates normalized counts.

**D)** Distribution of number of genes detected in nuclei in each cell type, by genotype. Blue, WT. Orange, QKI.
